# Thermodynamic, Electrochemical and Practical Constraints on Electromicrobial Formate Assimilation

**DOI:** 10.64898/2026.09.26.754500

**Authors:** Deniz Şinar, Razan Alharthi, Louis O. Anscombe, Shreya Singh, Farshid Salimijazi, Timothy J. Sheppard, David Specht, Buz Barstow

## Abstract

Electromicrobial production (EMP) technologies aim to combine renewable electricity, CO_2_, and engineered microbes to make energy-dense molecules at efficiencies exceeding photosynthesis. CO_2_ can be electrochemically reduced to formate, which is far easier to handle at the bench than H_2_ or an electrode, but formate carries only two electrons per carbon against the six in a biofuel. The remaining electrons must come from oxidizing additional formate, from H_2_ oxidation, or from extracellular electron uptake (EEU), and no rigorous comparison of these options coupled to the choice of carbon assimilation pathway currently exists. We calculate upper-limit efficiencies for butanol production by six carbon assimilation pathways, each paired with all three electron delivery mechanisms, using electrochemical parameters drawn from a survey of the recent literature. Electrical-to-butanol energy conversion efficiencies range from 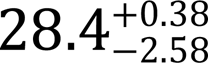 to 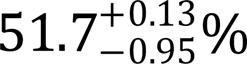, corresponding to solar-to-fuel efficiencies of 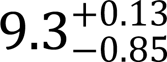 to 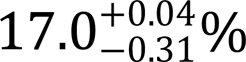, so even the least efficient route exceeds the 8% theoretical ceiling of algal photosynthesis. The serine variant of the reductive glycine pathway reaches an electrical energy conversion efficiency of 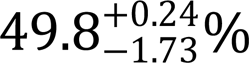 when using H_2_ oxidation, within 1.9 points of the most efficient pathway, and is the only high-efficiency option that tolerates O_2_. This makes an EMP system that combines electron delivery by formate coupled with the serine variant of the reductive glycine pathway highly attractive, as it presents few barriers to rapid, iterative engineering in the lab, and a high theoretical ceiling. Drawing both carbon and electrons from formate costs 6.2 points against H_2_ at a state-of-the-art whole-cell voltage (2.2 V). However, this small penalty is amplified three-fold by any rise in the CO_2_-to-formate cell voltage, and reaches 13.5 points at the highest whole-cell voltages reported for scaled-up CO_2_-to-formate electrolyzers, where formate-only operation falls to 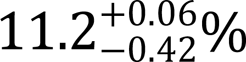 electrical-to-fuel and 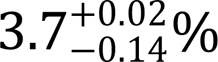 solar-to-fuel efficiency, below the ceiling of photosynthesis, against 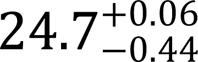 and 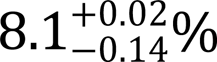 for H_2_ (only just above algal photosynthesis). Our choice between a formate-only system and one coupled to H_2_ oxidation or EEU therefore depends on our belief about the trajectory of CO_2_ reduction technology. If whole-cell voltages continue to fall at the rate of the past decade, formate alone is the right target, and the simplicity of its workflow is bought at low cost. However, if that improvement plateaus, the electron delivery mechanism must be swappable, and a system should be designed from the outset so that it can be. At the US Department of Energy SunShot target of 2¢ per kilowatt hour, the electricity to make a US gallon of butanol costs $1.40 for a formate-only system at the state of the art, rising to $5.45 at the highest scaled-up electrolyzer voltage reported, against $1.23 and $2.47 for formate and H_2_ systems.

## Introduction

Electromicrobial production (EMP) is a class of technologies that use microorganisms to convert renewable electricity and renewable carbon (typically atmospheric or captured CO_2_) into complex, energy-dense molecules with high efficiency^1,2^ (**Figure 1**). In principle, EMP could enable energy storage and the production of commodity chemicals, biofuels, food, and structural materials at low cost, high efficiency and high abundance, decoupling their supply from arable land, fossil carbon, or fermentable sugar^2–7^.

**Figure 1.**
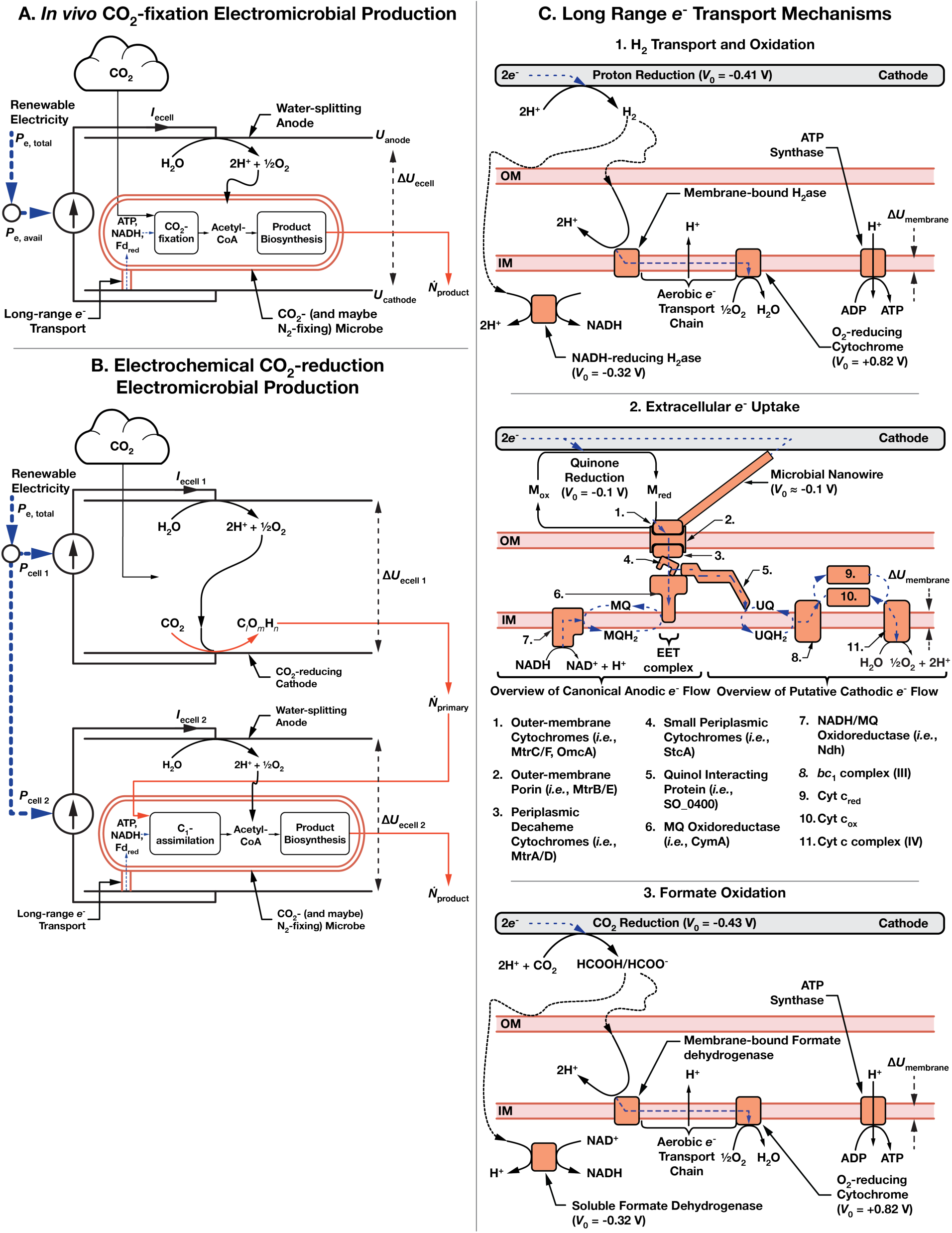
Schematic of electromicrobial production (EMP) systems that use electrochemically-reduced CO_2_. In this article, we consider EMP systems that utilize formate alone, but this framework can easily be extended to consider other CO_2_ reduction products, including carbon monoxide, methane, methanol, and acetate. (**A**) We consider *in vivo* CO_2_-fixation EMP systems using the Calvin-Benson-Bassham (CBB) cycle as points of comparison and to cross-check with earlier articles. (**B**) Production of C_1_ compounds involves an electrochemical system powered by renewable electricity that reduces atmospheric CO_2_. This C_1_ molecule is then transferred to a bio-electrochemical cell, where an engineered microbe converts it to butanol. (**C.1**) Microbial metabolism can be powered by H_2_ transport and oxidation, in which H_2_ is generated via electrochemical reduction on a cathode, transferred to the microbe by diffusion or stirring, and enzymatically oxidized to supply intracellular NAD(P)H and ATP; (**C.2**) by extracellular electron uptake (EEU), in which *e*^−^ are transferred along a microbial nanowire (part of a conductive biofilm) or by a reduced medium potential redox shuttle such as a quinone or flavin, and are then oxidized at the cell surface by the extracellular electron transfer (EET) complex. Electrons are then transported to the inner membrane, where reverse electron transport is used to regenerate NAD(P)H, reduced Ferredoxin (not shown), and ATP^2,13,14^; or (**C.3**) by transport and subsequent oxidation of formate, which is used to regenerate NAD(P)H, which can itself be oxidized to regenerate ATP. Parameters for these systems are shown in **Table 1**. This schematic is modified from earlier work on electromicrobial production^2,4–7^.

**Table 1.**
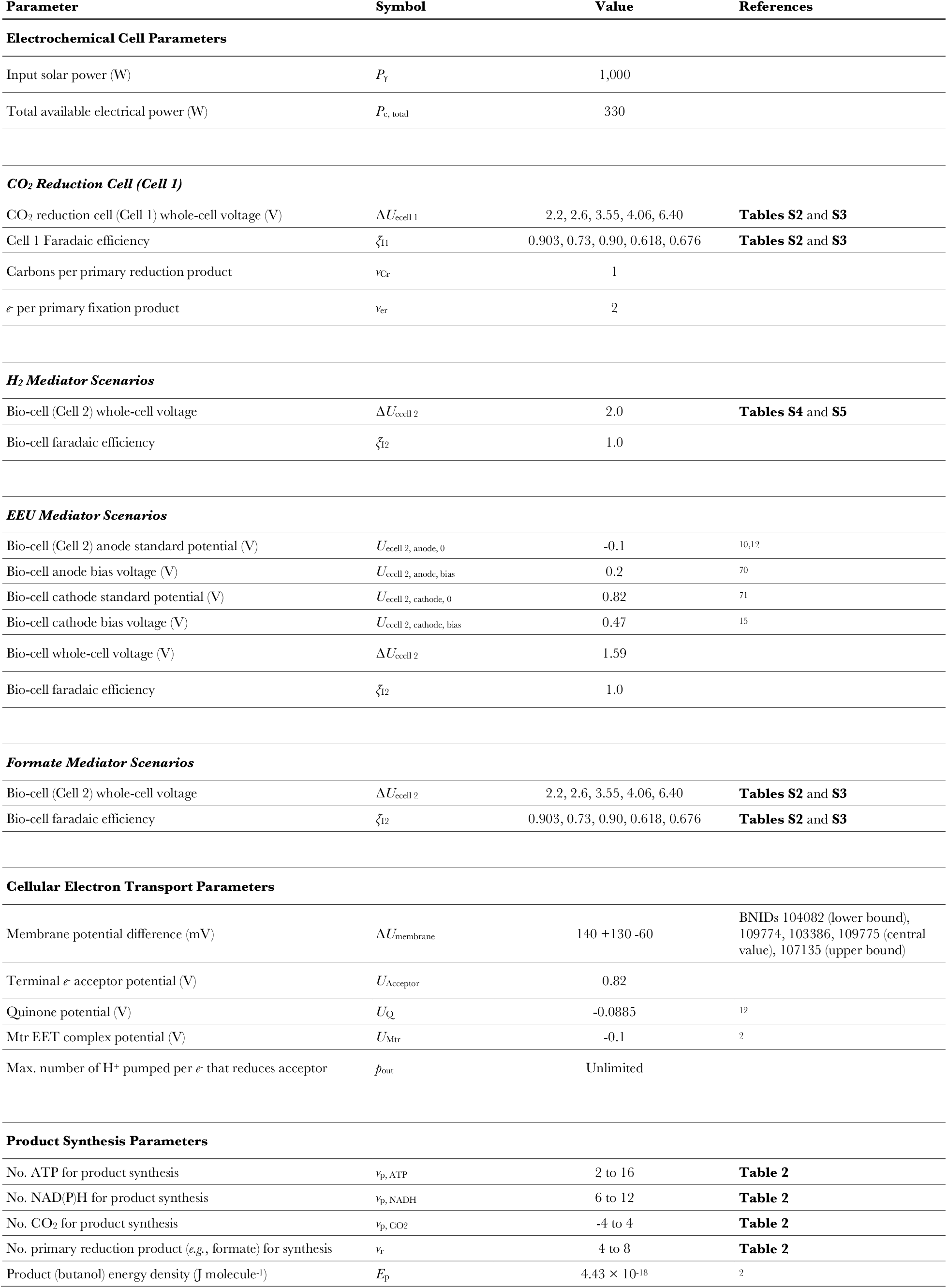
Model parameters for electromicrobial production of butanol from CO_2_ and formate. Model parameters used in this article are based upon parameters used in a previous analysis of the electromicrobial production of the biofuel butanol using *in vivo* CO_2_ fixation and a single formate assimilation pathway^2^. In this article, we have significantly updated the parameters of electrochemical formate and H_2_ production. A sensitivity analysis was performed for all key parameters in this work in an earlier article^2^.

| Parameter | Symbol | Value | References |
| --- | --- | --- | --- |
| **Electrochemical Cell Parameters** |  |  |  |
| Input solar power (W) | $P_{\gamma}$ | 1,000 | |
| Total available electrical power (W) | $P_{e, \text{total}}$ | 330 | |
| **CO<sub>2</sub> Reduction Cell (Cell 1)** |  |  |  |
| CO<sub>2</sub> reduction cell (Cell 1) whole-cell voltage (V) | $\Delta U_{\text{cell 1}}$ | 2.2, 2.6, 3.55, 4.06, 6.40 | **Tables S2 and S3** |
| Cell 1 Faradaic efficiency | $\zeta_{\text{f1}}$ | 0.903, 0.73, 0.90, 0.618, 0.676 | **Tables S2 and S3** |
| Carbons per primary reduction product | $\nu_{\text{Cr}}$ | 1 | |
| $e^-$ per primary fixation product | $\nu_{\text{er}}$ | 2 | |
| **H<sub>2</sub> Mediator Scenarios** |  |  |  |
| Bio-cell (Cell 2) whole-cell voltage | $\Delta U_{\text{cell 2}}$ | 2.0 | **Tables S4 and S5** |
| Bio-cell faradaic efficiency | $\zeta_{\text{f2}}$ | 1.0 | |
| **EEU Mediator Scenarios** |  |  |  |
| Bio-cell (Cell 2) anode standard potential (V) | $U_{\text{cell 2, anode, 0}}$ | -0.1 | 10,12 |
| Bio-cell anode bias voltage (V) | $U_{\text{cell 2, anode, bias}}$ | 0.2 | 70 |
| Bio-cell cathode standard potential (V) | $U_{\text{cell 2, cathode, 0}}$ | 0.82 | 71 |
| Bio-cell cathode bias voltage (V) | $U_{\text{cell 2, cathode, bias}}$ | 0.47 | 15 |
| Bio-cell whole-cell voltage (V) | $\Delta U_{\text{cell 2}}$ | 1.59 | |
| Bio-cell faradaic efficiency | $\zeta_{\text{f2}}$ | 1.0 | |
| **Formate Mediator Scenarios** |  |  |  |
| Bio-cell (Cell 2) whole-cell voltage | $\Delta U_{\text{cell 2}}$ | 2.2, 2.6, 3.55, 4.06, 6.40 | **Tables S2 and S3** |
| Bio-cell faradaic efficiency | $\zeta_{\text{f2}}$ | 0.903, 0.73, 0.90, 0.618, 0.676 | **Tables S2 and S3** |
| **Cellular Electron Transport Parameters** |  |  |  |
| Membrane potential difference (mV) | $\Delta U_{\text{membrane}}$ | 140 +130 -60 | BNIDs 104082 (lower bound), 109774, 103386, 109775 (central value), 107135 (upper bound) |
| Terminal $e^-$ acceptor potential (V) | $U_{\text{Acceptor}}$ | 0.82 | |
| Quinone potential (V) | $U_{\text{Q}}$ | -0.0885 | 12 |
| Mtr EET complex potential (V) | $U_{\text{Mtr}}$ | -0.1 | 2 |
| Max. number of H<sup>+</sup> pumped per $e^-$ that reduces acceptor | $p_{\text{out}}$ | Unlimited | |
| **Product Synthesis Parameters** |  |  |  |
| No. ATP for product synthesis | $\nu_{\text{p, ATP}}$ | 2 to 16 | **Table 2** |
| No. NAD(P)H for product synthesis | $\nu_{\text{p, NADH}}$ | 6 to 12 | **Table 2** |
| No. CO<sub>2</sub> for product synthesis | $\nu_{\text{p, CO2}}$ | -4 to 4 | **Table 2** |
| No. primary reduction product (*e.g.*, formate) for synthesis | $\nu_{\text{r}}$ | 4 to 8 | **Table 2** |
| Product (butanol) energy density (J molecule<sup>-1</sup>) | $E_{\text{p}}$ | $4.43 \times 10^{-18}$ | 2 |

The term electromicrobial production covers any system in which the carbon and the reducing power for microbial biosynthesis both originate from CO_2_ and renewable electricity. In conventional fermentation, carbon and electrons arrive together in a sugar molecule whose carbon was fixed by photosynthesis and whose electrons came from water splitting. In an EMP system, they may also arrive together, or separately, but they originate from an electrolyzer and an atmospheric or captured CO_2_ stream (**Figures 1A** and **1B**).

Several mechanisms are available for delivering electricity into metabolism, and the choice among them should be a central design decision (**Figure 1C**). Electrons can be supplied by oxidation of electrochemically generated H_2_ (**Figure 1C.1**); by extracellular electron uptake (EEU) from a cathode, through conductive appendages or soluble redox mediators (**Figure 1C.2**); or by oxidation of an electrochemically reduced carbon compound like formate, which supplies carbon and reducing power in the same molecule (**Figure 1C.3**). The term microbial electrosynthesis is often used for the EEU case specifically^8,9^, though the boundary is not sharp: H_2_-mediated systems fall inside or outside it depending on whether H_2_ production is biologically enhanced at the cathode^2^. We use electromicrobial production for the general case^1^, so that systems using different electron delivery mechanisms can be compared on the same footing.

The three electron delivery mechanisms considered in this article differ in a way that is easy to overlook. Electrons must reach metabolism at the potential of NAD(P)H (−320 mV vs. Standard Hydrogen Electrode (SHE)) or reduced ferredoxin (≈ −420 mV vs. SHE) before they can be used to reduce carbon. H_2_ oxidation delivers electrons at −420 mV and formate oxidation at −430 mV, close enough that little energy is wasted. In systems mediated by EEU, electrons are delivered in the potential window of the outer membrane conduit, from ≈ +100 to −100 mV vs. SHE^10^, far too high to be useful for NAD(P)H or ferredoxin regeneration. These electrons must be raised to the potential of NAD(P)H or ferredoxin by reverse electron transport^2,11–14^. The efficiency impact of this up-conversion was not obvious from initial inspection.

The thermodynamic case for EMP has been established for over a decade. Over the past several years, our group has developed a theoretical framework for calculating the upper-limit efficiency of EMP systems^2,4–7^. In these works, we demonstrated that the upper-limit efficiency of EMP could far exceed the theoretical upper limit efficiency of all known forms of photosynthesis. This framework has focused primarily on the choice of electron delivery mechanism, H_2_ oxidation or extracellular electron uptake (EEU), and, more recently, on the choice of target product. Similarly, Liu *et al*.^15^ demonstrated that a system combining the H_2_-oxidizing microbe *Cupriavidus necator* (also known as *Ralstonia eutropha*) and a proton-reducing electrode system capable of operating at neutral pH (the Bionic Leaf) could produce the biofuel isopropanol with an efficiency far exceeding that of photosynthesis if supplied with even a commercially available solar photovoltaic. Likewise, Leger *et al*. calculated that EMP could produce proteins at a cost far below that of conventional agriculture^3^.

In practice, EMP has already begun to leave the lab. Solar Foods has operated a commercial-scale facility producing single-cell protein from CO_2_ and H_2_ since 2024^16,17^, and Circe Bioscience opened a pilot plant producing triglycerides from the same inputs^18,19^. However, EMP has not yet achieved its original promise of delivering highly abundant commodity chemicals at ultra-low-cost^1,20^. Both Solar Foods and Circe’s systems use H_2_-oxidizing chemolithoautotrophs running their native Calvin cycle, the one route to EMP that requires neither a new carbon fixation pathway nor a new chassis organism^19,21^. However, while H_2_-oxidizing systems are relatively easy to set up and demonstrate in the lab because organisms naturally oxidize H_2_ and fix CO_2_, they face a major scale-up challenge due to H2’s insolubility in water. Counter-intuitively, this challenge is actually greatest at small scale ^2^. At very large scale – systems scaled to accept power at a rate of 1 MW or greater – the fraction of input power required for agitation becomes negligible^2^. However, at smaller scales, precisely the scale where a startup would operate, this power draw poses a significant efficiency and cost burden. While this may be tolerable when producing a high-value product like a protein – both companies targeted high-value products ^17,18^ – it is not tolerable when producing commodity chemicals like biofuels. Furthermore, conventional hydrogen electrolyzers operate at high temperature, adding a further energetic burden that is not obvious at lab scale, nor accounted for in our original thermodynamic analysis^2^. The cobalt-based electrode pair used in the Bionic Leaf, a cobalt phosphate anode^22^ and a cobalt-phosphorus alloy cathode^15^, circumvents these problems by operating at ambient temperature and neutral pH, and *in situ* – removing the need to heavily agitate^23^. However, Liu *et al*. only reported 384 hours of stable operation for the cobalt-phosphate electrode system in a bioelectrochemical system^15^. By comparison, recent CO_2_-to-formate electrolyzers report 1,000 to 8,000 hours^24,25^ of operation, while commercial alkaline and PEM (Proton Exchange Membrane) systems have lifespans of years. The cobalt-phosphate chemistry underlying the Bionic Leaf anode was commercialized by Sun Catalytix, which subsequently redirected its efforts to flow batteries and whose assets were acquired by Lockheed Martin in 2014^26^. The chemistry was not deployed at scale for water splitting.

Extracellular electron uptake could allow us to circumvent the H_2_ solubility problem. Electrons reach the cell either through a solid matrix, typically a conductive appendage synthesized by the microbes like a nanowire^27^ or cable^28^, or through a soluble redox mediator like anthrahydroquinone-2,6-disulfonate (AHDS, or AQDS_red_^14^). In the solid matrix case, the energy losses due to electron transport come from ohmic losses along the nanowires or cables^2^. The upper bound on performance appears to be high: cable bacteria of the family *Desulfobulbaceae* produce filaments up to a centimeter in length with conductivities approaching those of metals^28^, indicating that ohmic losses in long-range biological electron transport need not be significant^2^. Furthermore, having the microbes embedded in a solid matrix potentially dramatically simplifies product separation. In the mediated case the shuttle is much more soluble than H_2_, which removes much of the mass transfer problem that H_2_ presents. Furthermore, new mediators have recently become available that address some of the solubility issues of AQDS_red_^29^. In either case the whole-cell voltage of an EEU bioelectrochemical cell can be as low as 1.59 V, against 2.0 V for a H_2_-producing cell (**Table 1**), although the reverse electron transport needed to raise electrons from the extracellular electron transfer complex to the potential of NAD(P)H consumes some of that advantage.

The difficulty with EEU is not thermodynamic but in engineering. The organisms in which EEU is best characterized are almost exclusively inconvenient to work with. *Shewanella oneidensis*, in which the electron uptake pathway has been most thoroughly characterized^13,14^, is genetically tractable but grows slowly, and the cable bacteria at the high end of the conductivity range are unlikely to be amenable to genetic manipulation at all ^30^.

Native electron uptake capability may occur in more tractable organisms. *Vibrio natriegens*, possibly the most engineerable microorganism known^31^, possesses a characterized Mtr-type extracellular electron transfer (EET) conduit consisting of homologs of CymA, PdsA and MtrCAB^32^, and respires anodically both directly and through artificial mediators^33^. These demonstrations are of electron export rather than uptake, and the currents involved are modest. Whether such a conduit can be driven in reverse at a useful rate, as it can in *S. oneidensis*^13,14^, is not yet established, although this seems highly likely.

Even so, EEU remains a slow and demanding route to iterate. Where the solid matrix is used, every cycle of engineering is gated by the time needed to establish a biofilm on an electrode, which takes days to weeks against hours for a planktonic culture.

Soluble mediators that interface with the EEU complex relieve the biofilm growth constraint during initial engineering, but they impose one of their own. H_2_ and formate are consumed: oxidizing formate yields CO_2_, which leaves the medium as a gas, and oxidizing H_2_ yields protons, most of which are incorporated into the reduced product. Neither mediator leaves much behind to perturb the medium. On the other hand, soluble mediators that interface with the EEU complex are not consumed but converted. Oxidized AQDS remains in the reactor at the same concentration, where it is potentially toxic to the cells and cannot be diluted away. Simply adding more mediator is not an option, both because the mediator is used at near its solubility limit and because accumulating degradation products are likely to be worse for the culture than the mediator itself. The mediator must therefore be continuously re-reduced electrochemically, which means the electrochemical cell is part of every experiment, not just part of the eventual process.

The conditions required for EEU-mediated EMP are technically demanding. Microbes that take up electrons in nature do so under constraints that differ by lifestyle. Neutrophilic chemolithotrophic iron oxidizers are microaerophiles, requiring opposing gradients of oxygen and iron: oxygen serves as the terminal electron acceptor for the proton gradient that drives reverse electron transport, but in excess it allows abiotic oxidation to outpace the biological reaction^34^. The insoluble ferric oxyhydroxides produced by either route would otherwise encase the cells, and are managed by the excreted stalks of *Gallionella* and the tubular sheaths of *Leptothrix*, which coordinate precipitation away from the cell surface^34^. *Acidithiobacillus ferrooxidans* mitigates both problems by operating at pH 1–2, where abiotic oxidation of Fe^2+^ is slow and dissolved ferrous iron can reach 10^-1^ M, some sixteen orders of magnitude above circumneutral environments, and where the ferric product is substantially more soluble^35^. Anoxygenic phototrophs like *Rhodopseudomonas palustris* avoid the competition entirely by operating anaerobically, using light rather than a terminal oxidase to raise the energy of electrons taken from Fe^2+^ (Ref. ^36^).

An electrode presents a combination of constraints on EMP-mediated EEU matching none of those seen in nature. There is no iron, so the precipitation problem disappears, but oxygen reduction at the cathode is a competing reaction. Oxygen reduction on carbon is slow, requiring approximately 300 mV of overpotential to initiate^37^, and proceeds largely by the two-electron route to hydrogen peroxide rather than the four-electron route to water^38^. Activity is further suppressed at neutral pH: undoped oxygen-rich carbon electrodes produce a measurable peroxide current only at 0.266 V vs. RHE in neutral electrolyte^25^ (≈ −0.15 V vs. SHE at pH 7), which is at or below the potential at which EEU systems operate. Some peroxide production is therefore likely at the more reducing end of the EEU range, though the rates involved are likely modest. Because this reaction is kinetically rather than transport limited, electrode composition offers a route to mitigation with no counterpart in the precipitation problem faced by neutrophilic iron oxidizers: onset potentials and two-electron selectivity on carbon can be shifted by several hundred millivolts by doping with Lewis acid sites or metal clusters ^25,37^.

Management of oxygen is likely a key constraint on EMP-mediated EEU. An oxygen gradient near the electrode is nonetheless unavoidable, since the cells must sit where the electrons are while also requiring oxygen for the terminal oxidase. The oxygen concentration that is optimal at the electrode surface is therefore probably not the concentration that is optimal in the bulk medium, and the difference between them depends on current density, mixing, and the thickness of the biofilm. Whether the resulting constraint on oxygen concentration is tighter or looser than in the neutrophilic iron-oxidizing case is not clear, and the uptake pathway has likely never been under selection to function at high oxygen concentration.

Irrespective of whether EEU operates with a solid matrix or a soluble mediator, the experimental workflow requires a bioelectrochemical reactor, likely with atmospheric control, rather than a flask. The additional apparatus and expertise this demands presents a real barrier for most students and postdocs. A route with a high efficiency ceiling but a design-build-test cycle measured in weeks, and an apparatus that requires deep skills in both genetics and electrochemistry, a combination few people have, is not obviously preferable to one with a lower ceiling that can be iterated in days on the bench. This is not an argument against EEU. Upper-limit efficiency matters profoundly, but it is not the only thing that does, and we have come to believe that the choice between electron delivery mechanisms cannot be made on efficiency alone.

Formate avoids both sets of problems. Formic acid and its conjugate base are fully miscible with water, so much less energy needs to be spent on agitation to bring the mediator into contact with the cells. Because the mediator is soluble and stable, the CO_2_-reducing cell no longer needs to share a compartment, or a set of operating conditions, with the culture. The electrolyzer can be optimized as an electrolyzer and the bioreactor as a bioreactor. CO_2_-to-formate electrolysis runs at ambient temperature, and in some configurations requires active cooling rather than heating^39^, so the thermal burden carried by conventional hydrogen electrolyzers does not arise. State-of-the-art whole-cell voltages for CO_2_ reduction to formate have fallen from around 3.5 V to 2.2 V over the past decade at faradaic efficiencies above 90% (**Table S2**), narrowing the gap with H_2_ considerably.

Formate is also, by some distance, the easiest of the three mediators to work with at the bench. It is bought or made, stored, and measured into a flask like any other medium component. When it is oxidized the carbon simply leaves as CO_2_. The experiment does not require an electrode or gas handling. This means that the design-build-test cycle for formate-mediated EMP runs at the speed of the organism rather than the speed of the apparatus, which matters a great deal when the engineering task ahead requires so many cycles.

The current state of development of CO_2_ to formate reduction means that formate-mediated EMP carries a cost that the other mediators do not. Formate is only lightly reduced, carrying two electrons per carbon, whereas a biofuel like butanol carries six. Formate can supply all of the carbon a cell needs to make something like butanol but only a third of the electrons, and the balance must come from somewhere. If that balance is supplied by oxidizing additional formate, then roughly two-thirds of the formate entering the cell is oxidized back to CO_2_ in order to reduce the remaining third to product, and the electrochemical cell must therefore reduce three molecules of CO_2_ for every carbon that reaches the product. This means that any improvement or degradation in the CO_2_ to formate reduction voltage is amplified threefold. Alternatively, the additional electrons can be supplied by H_2_-oxidation or by EEU, which given today’s state of the art recovers efficiency at the cost of reintroducing the constraints that made formate attractive in the first place.

Selecting the right mediator to pair with formate-assimilation today and in the future requires a comparison that does not currently exist. The efficiency of an EMP system depends jointly on the electron delivery mechanism and on the carbon assimilation pathway used *in vivo*. A pathway that is expensive in ATP is penalized more heavily by a mediator that delivers electrons at a higher potential, since fewer protons can be pumped per electron to regenerate it; a pathway that releases CO_2_ as an intermediate wastes electrons that were paid for at the electrolyzer. Bar-Even and colleagues mapped the space of natural and synthetic formate assimilation pathways in detail and assessed them for engineering tractability, using proxies like overlap with central metabolism and the oxygen sensitivity of the constituent enzymes^40–42^. Separately, Claassens *et al*. surveyed experimental growth data to rank electron mediators by the energetic efficiency of feedstock bioconversion^1^. What has not been attempted is a thermodynamic ranking of the carbon assimilation pathways themselves, crossed against the choice of mediator, for the upper-limit efficiency with which electricity is converted into product. Meanwhile, the electrochemical parameters on which any such ranking depends have moved substantially: the whole-cell voltages we used in our own earlier analysis^2^ are no longer state of the art.

The impact of electron and carbon delivery choice on the energy conversion efficiency from electricity (or sunlight) to product is not obvious by inspection. Efficiency is far from being the only selection criterion that matters, but it is the only one that can be calculated before any strain is built, and it sets the performance ceiling of the system. That ceiling is what determines whether an EMP system can beat photosynthesis, which is the point of building one. Absent that calculation, the choice of chassis and pathway is made on other grounds, and those grounds are not necessarily correlated with where the ceiling lies. This article provides that calculation.

## Theory

In this article, we extend a computational framework developed in our previous works^2,7^ to calculate the upper limit efficiencies of theoretical EMP systems that convert either CO_2_ or the C_1_ molecule formate into the biofuel butanol. Butanol was selected for its compatibility with modern engine technologies^43,44^; its relatively simple biochemical synthesis pathway, and its successful production by engineered microorganisms^45,46^.

Symbols appearing in this article are listed in **Table S1**. Model parameters with corresponding values are listed in **Table 1**, and a representative selection of state-of-the-art and industrial electrochemical voltages for CO_2_ reduction to formate, and proton reduction to H_2_ are shown in **Tables S2** to **S5**. We restate elements of this theory from earlier articles^2,7^ here for the convenience of the reader. The derivation for these calculations can be found in the supplemental material of our previous works by Salimijazi *et al.*^2^ and Wise *et al*.^7^.

We consider a bio-electrochemical system with access to an unlimited reservoir of atmospheric CO_2_ for *in vivo* CO_2_-fixation through the well-known Calvin-Benson-Bassham cycle (**Figure 1A**) as a control, or electrochemical reduction into formate that will be fed into microbial metabolism (**Figure 1B**). Electrons are supplied to microbial metabolism for reduction reactions through oxidation of electrochemically-produced H_2_ (**Figure 1C.1**), Extracellular Electron Uptake (EEU) (**Figure 1C.2**), or oxidation of electrochemically-produced formate (**Figure 1C.3**). In all three electron delivery mechanisms, the cathode provides reducing power allowing regeneration of NAD(P)H, reduced ferredoxin (Fd_red_), and ATP. Microbial maintenance energy is assumed to be negligible^2^, thus almost all electrical energy is put toward producing butanol as the target molecule with an internal energy *E*_fuel_, at a rate of *Ṅ*_fuel_. Reaction diagrams and full reaction lists for the CO_2_-fixing Calvin-Benson-Bassham (CBB) cycle; glyceraldehyde variant of the formolase formate-assimilation pathway^42^; dihydroxyacetone (DHA) variant of the formolase pathway^42^; glycine-reductase variant of the reductive glycine pathway^42^; serine-variant of the reductive glycine pathway^42^; and Wood-Ljungdahl (WL) formate-assimilation pathway are shown in **Figures 2** to **7** and **Datasets S1A** to **S1F**. The synthesis pathway for butanol is shown in **Figure 8** and **Dataset S1G**.

**Figure 2.**
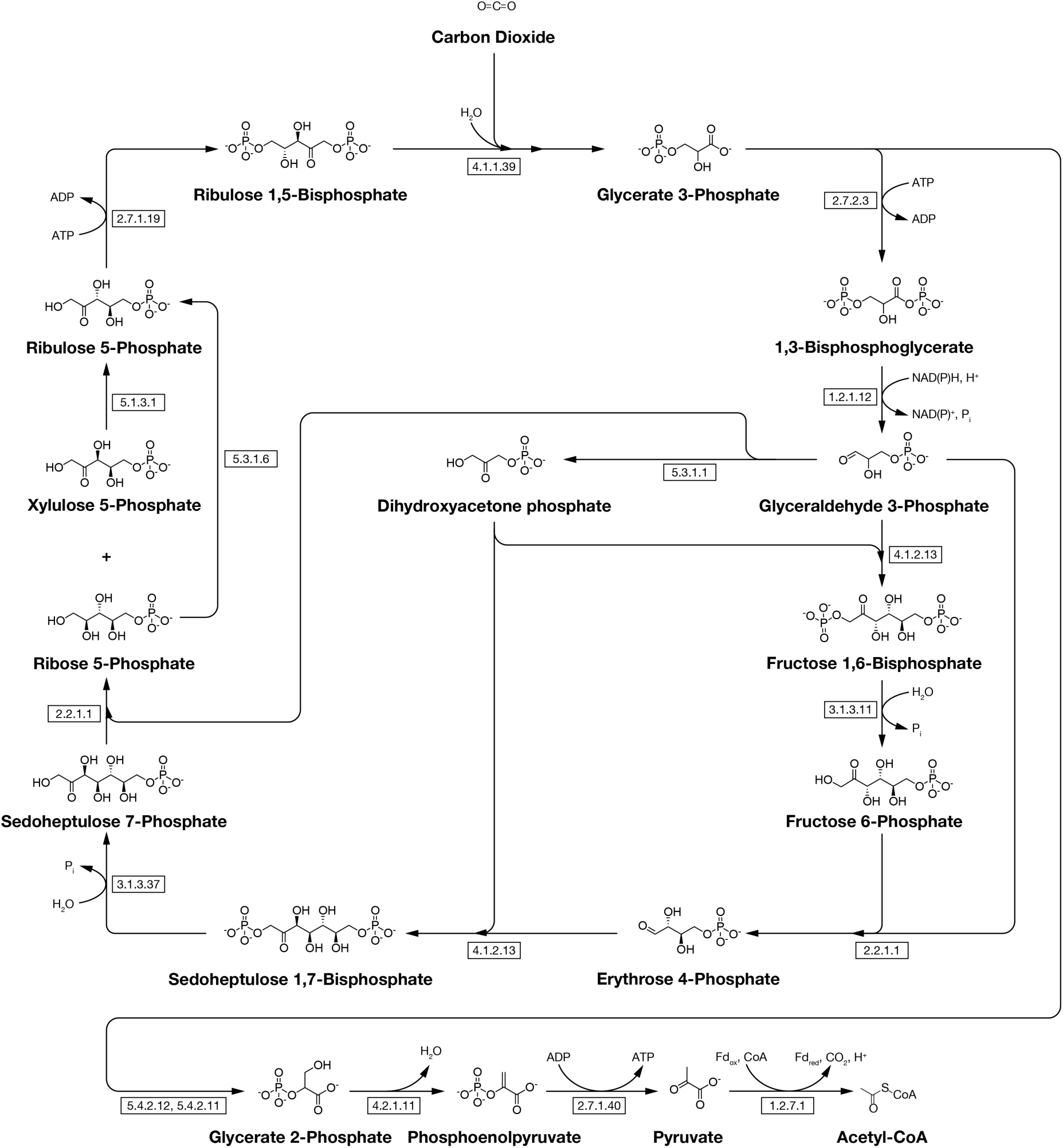
The Calvin-Benson-Bassham (CBB) or Calvin cycle. The Calvin cycle is the predominant method of carbon- fixation on earth, present in plants, algae, and cyanobacteria. This article considers two variants of the Calvin cycle: in the first (P1a-CBB) carbon is sourced directly from the atmosphere by enzymatic fixation by RuBisCO (E.C. 4.1.1.39); in the second (P1b-FDH-CBB) carbon is supplied as formate and then oxidized to CO_2_ by formate dehydrogenase (FDH). As with all CO_2_-fixation and formate assimilation pathways considered in this article, we present the pathway up until the production of acetyl-CoA. The pathway for production of butanol from acetyl-CoA is shown in Figure 8. Full details of the reactions shown here can be found in **Dataset S1A**. ATP, NAD(P)H, CO_2,_ and HCO_2_^-^ requirements for these pathways are calculated by P1_A_-CBB._PY_ and P1_B_-FDH-CBB._PY_ in the GitHub repository for this article^47^.

The energy conversion efficiency of electricity to butanol, *η_EF_*, is calculated from the ratio of the amount of chemical energy stored per second (*Ṅ*_fuel_ *E*_fuel_), relative to the power input to the system, *P*_e, total,_

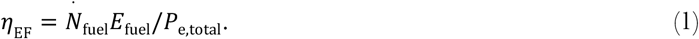

_·_Additionally, the energy required to generate one mole of desired hydrocarbon product, *L*_EF_, is calculated from the following,

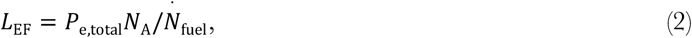

where *N*_A_ is Avogadro’s number.

For the case of *in vivo* CO_2_ fixation (**Figure 1A**) as our control, the upper limit electrical-to-chemical conversion efficiency is calculated using the energy density of butanol relative to the charge required to synthesize it from CO_2_ (*eν*_ef_; the fundamental charge multiplied by the number of electrons required for synthesis) and the potential difference across the bio-electrochemical cell (Δ*U*_ecell_)^2^,

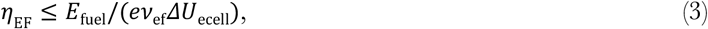

Therefore, the input electricity required for a mole of butanol is,

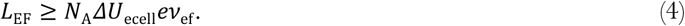

The focus of this article is on systems in which CO_2_ is electrochemically reduced to formate. As we noted earlier, formate is only lightly reduced, carrying only 2*e*^−^ per carbon. Thus, formate can provide the carbon and some electrons necessary for butanol biosynthesis, but the remainder of electrons needed for this process must be supplied by H_2_ oxidation, EEU, or oxidation of more formate. The upper limit electricity-to-product conversion efficiency is calculated as

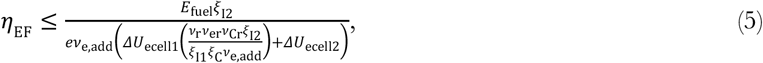

where *ν*_e, add_ represents the number of electrons required to fully reduce formate to the final product (*i.e.*, butanol), Δ*U*_ecell1_ is the potential difference across the CO_2_-reducing cell, *ν*_r_ is the required number of primary reduction products (*i.e.*, formate) needed to generate a final product, *ν*_er_ is the number of electrons to reduce CO_2_ to this primary product (*e.g.*, 2 in the case of formate), *ν*_Cr_ is the number of carbon atoms per primary fixation product (*i.e.*, 1 in the case of formate), *ξ*_I2_ is the faradaic efficiency of the bio-electrochemical cell, *ξ*_I1_ is the faradaic efficiency of the primary abiotic cell 1, and *ξ*_C_ is the carbon transfer efficiency from electrochemical cell 1 to cell 2^2^.

Therefore, the electrical energy required to produce one mole of 1-butanol is calculated as

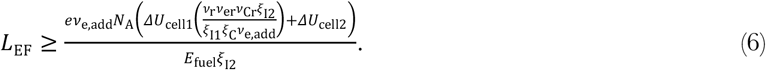

The number of electrons needed for fuel synthesis (*ν*_ef_ or *ν*_e, add_) is calculated from a model of electron uptake by H_2_-oxidation or EEU (first introduced in Salimijazi *et al*.^2^, and experimentally suggested by Rowe *et al*.^13,14^), and from the number of Fd_red_, ATP, and NAD(P)H required by the metabolic pathway used for fuel biosynthesis^2^ (*ν*_f, Fd_, *ν*_f, ATP_, and *ν*_f, NADH_). For computations, we assume NADH and NADPH as interchangeable given their identical redox potentials. For electron delivery by H_2_,

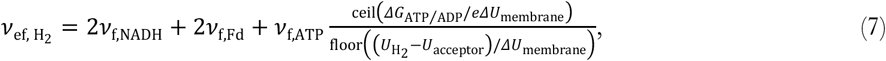

where Δ*G*_ATP/ADP_ is the free energy required for the regeneration of ATP, Δ*U*_membrane_ is the potential difference across the cell’s inner membrane due to the proton gradient, *U*_H2_ is the standard redox potential of proton reduction to H_2_, *U*_acceptor_ is the potential of the electron acceptor, *U*_NADH_ is the potential of NADH, and *U*_Fd_ is the potential of ferredoxin. The ceil function rounds up to the nearest integer, while the floor function rounds down to the nearest integer.

Furthermore, for electron delivery by EEU,

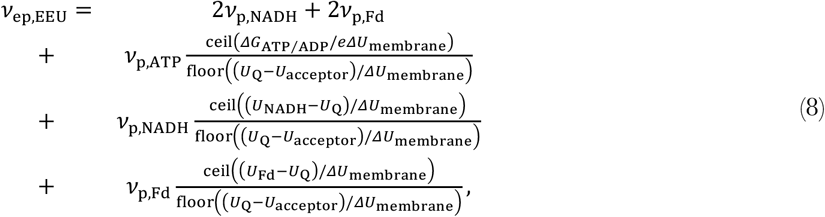

where *U*_Q_ is the redox potential of the quinone inner membrane electron carrier.

The set of reactions for product synthesis are balanced in a custom flux balance analysis code that calculates the required NAD(P)H, ATP, CO_2_, and formate inputs needed to produce a single target molecule^7^. For computational analysis, a stoichiometric matrix is generated wherein the vector ***ṅ*** encodes the change in number of molecules over a single round of the reaction cycle; **S_p_** is a matrix that encodes the stoichiometry for each reaction in the pathway; and ***ν*** is a flux vector describing the number of times each reaction is utilized throughout a reaction cycle,

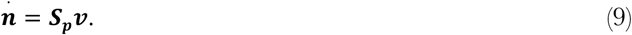

In this framework, *ṅ_i_* is constrained to be zero for each reactant designated as an intermediate (*e.g.*, acetyl-CoA, acetoacetyl-CoA, *etc.*; but not CO_2_, NAD(P)H, or ATP) in the reaction network (*i.e.*, its number should not grow or shrink over an entire reaction cycle, leaving the host cell’s chemical state unchanged over time). The optimized stoichiometry for synthesis of butanol by each pathway considered in this article is available in the code repository for this article^47^. Calculations for the CBB cycle were used as a control by running reactions through the program sequence and comparing them to pre-calculated values for ATP, NADH, and CO_2_ input requirements.

As a sanity check, the total electron demand computed for each pathway must equal 24 *e*^-^ per butanol when carbon is supplied as CO_2_, or 16 additional *e*^-^ when four formate molecules supply the carbon and 8 of the required electrons. All pathways satisfy this budget.

A second program^2^ then uses values for the required NAD(P)H, Fd_red_, and ATP (**Table 2**) to produce butanol in each pathway and calculates the required input energy necessary for biosynthesis based on the energy density and molecular weight of butanol.

**Table 2.** Overview of CO_2_-fixation and formate-assimilation pathways. Net molecular input requirements for butanol synthesis by the Calvin-Benson-Bassham CO_2_-fixation cycle and formate assimilation pathways. All input requirements are for a single butanol molecule. The 7^th^ column refers to the figure showing the reactions unique to each pathway that convert CO_2_ or formate to acetyl-CoA. The 8^th^ column refers to a dataset with further reaction information. The shared reactions that synthesize butanol from acetyl-CoA are shown in **Figure 8** and further information is detailed in **Dataset S1G**. The molecular input requirements are calculated by scripts in the _BALANCE_ subfolder of the GitHub repository associated with this article^47^.

| Pathway | Pathway Name | ATP | NAD(P)H | HCO<sub>2</sub><sup>-</sup> | CO<sub>2</sub> | Reaction Figure | Reaction Dataset |
| --- | --- | --- | --- | --- | --- | --- | --- |
| P1a-CBB | Calvin-Benson-Bassham cycle | 14 | 12 | 0 | 4 | **2** | **S1A** |
| P1b-FDH-CBB | Calvin-Benson-Bassham cycle supplied by formate dehydrogenase | 14 | 8 | 4 | 0 | **2** | **S1A** |
| P2a-Form-Glyc | Formolase pathway, glyceraldehyde variant, no CO<sub>2</sub> recycling | 16 | 8 | 8 | -4 | **3** | **S1B** |
| P2b-Form-Glyc-R | Formolase pathway, glyceraldehyde variant, CO<sub>2</sub> recycling | 16 | 12 | 4 | 0 | **3** | **S1B** |
| P3a-Form-DHA | Formolase pathway, dihydroxyacetone variant, no CO<sub>2</sub> recycling | 12 | 6 | 6 | -2 | **4** | **S1C** |
| P3b-Form-DHA-R | Formolase pathway, dihydroxyacetone variant, CO<sub>2</sub> recycling | 12 | 8 | 4 | 0 | **4** | **S1C** |
| P4-rGly-Gly | Reductive glycine pathway, glycine reductase variant | 4 | 8 | 4 | 0 | **5** | **S1D** |
| P5-rGly-Ser | Reductive glycine pathway, serine variant | 4 | 8 | 4 | 0 | **6** | **S1E** |
| P6-FDH-WL | Wood-Ljungdahl pathway supplied by formate dehydrogenase | 2 | 8 | 4 | 0 | **7** | **S1F** |

## Results and Discussion

In this article we calculate the upper-limit efficiency of butanol production for six carbon assimilation pathways, each coupled to three electron delivery mechanisms, using electrochemical parameters drawn from a current survey of the CO_2_-to-formate and H_2_ production literature (**Tables S2** to **S5**).

We consider the Calvin-Benson-Bassham (CBB) cycle (**Figure 2**)^48–50^, which can fix atmospheric CO_2_ directly or assimilate formate depending on whether formate dehydrogenase is used to supply CO_2_ and NADH or NADH alone; the glyceraldehyde variant of the formolase pathway^42^ (Form-Glyc; **Figure 3**); the dihydroxyacetone variant of the formolase pathway^42^ (Form-DHA; **Figure 4**); the glycine reductase variant of the reductive glycine pathway^42^ (rGly-Gly; **Figure 5**); the serine variant of the reductive glycine pathway^42^ (rGly-Ser; **Figure 6**); and the formate dehydrogenase variant of the Wood-Ljungdahl pathway, also known as the reductive acetyl-CoA pathway^51^ (FDH-WL; **Figure 7**).

**Figure 3.**
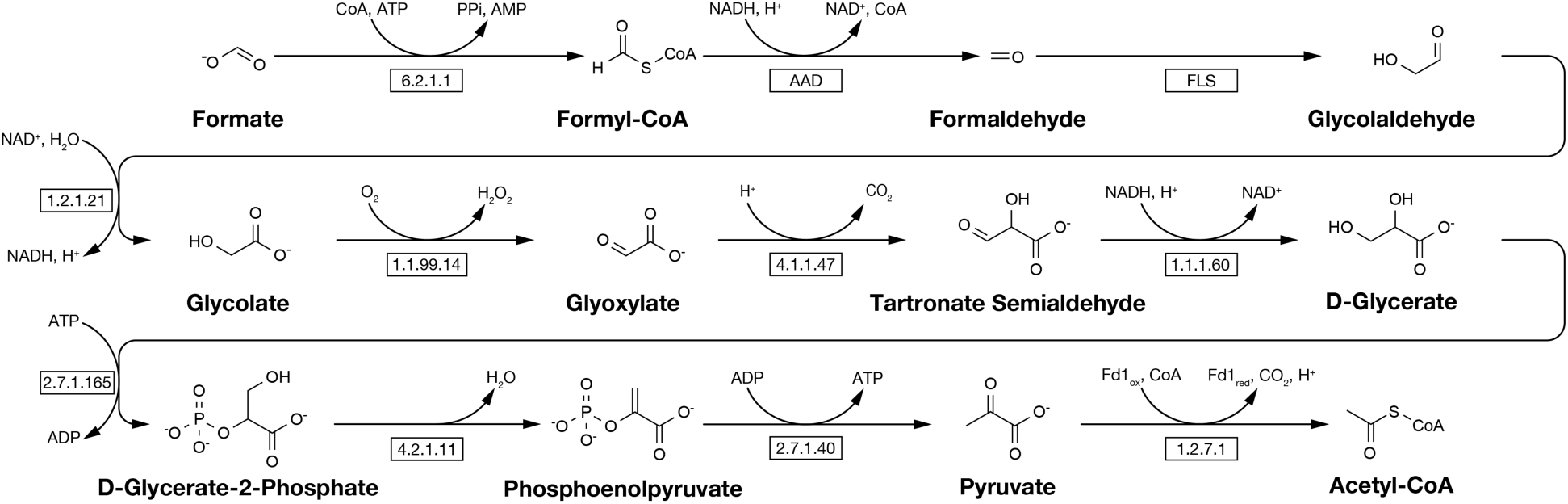
Glyceraldehyde variant of the formolase pathway (Form-Glyc). Siegel *et al*.^68^ computationally designed the enzyme formolase for use in novel synthetic formate assimilation pathways. Full details of the reactions shown here can be found in **Dataset S1B**. Note that CO_2_ is released during the conversion of pyruvate to acetyl-CoA. We consider a variant of this pathway where CO_2_ is recycled back to formate by running formate dehydrogenase in reverse at the expense of a single NADH. ATP, NAD(P)H, CO_2_ and HCO_2_^-^ requirements for these pathways are calculated by P2_A_-F_ORM_-G_LYC_._PY_ and P2_B_-F_ORM_-G_LYC_-R._PY_ in the GitHub repository for this article^47^.

**Figure 4.**
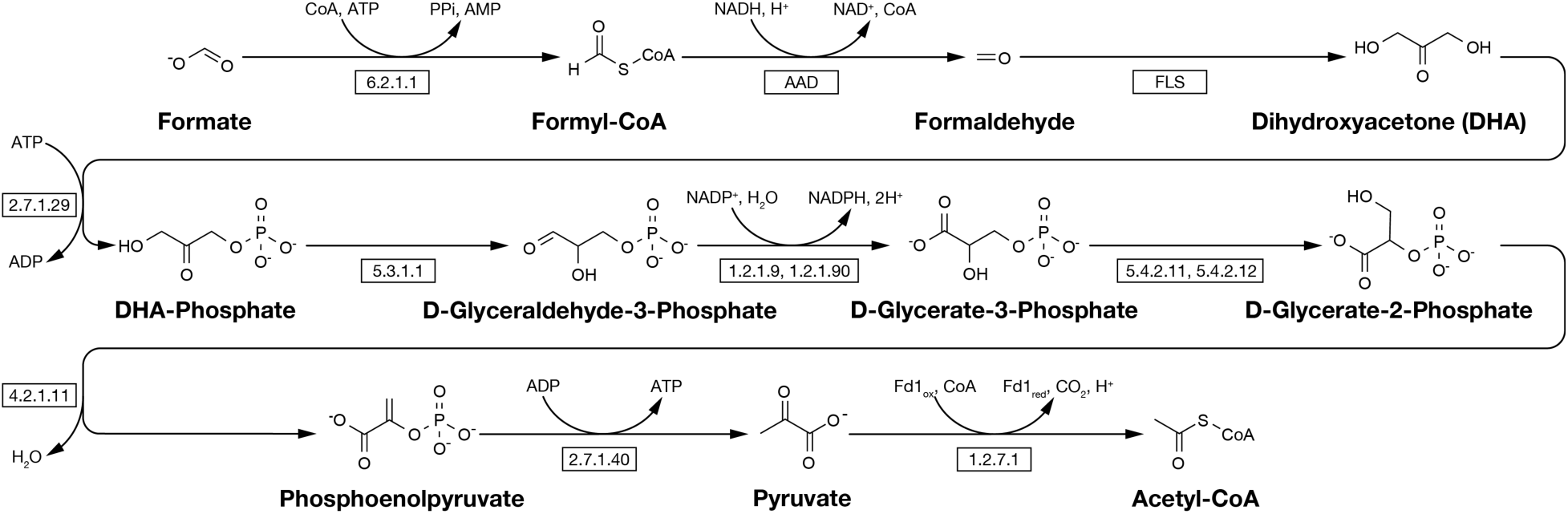
Dihydroxyacetone variant of the formolase pathway (Form-DHA). Siegel *et al*.^68^ computationally designed the enzyme formolase for use in novel synthetic formate assimilation pathways. The dihydroxyacetone variant^42^ functions at high formaldehyde concentrations. Full details of the reactions shown here can be found in **Dataset S1C**. Note that CO_2_ is released during the conversion of pyruvate to acetyl-CoA. We consider a variant of this pathway where CO_2_ is recycled back to formate by running formate dehydrogenase in reverse at the expense of a single NADH. ATP, NAD(P)H, CO_2_ and HCO_2_^-^ requirements for these pathways are calculated by P3_A_-F_ORM_-DHA._PY_ and P3_B_-F_ORM_-DHA-R._PY_ in the GitHub repository for this article^47^.

**Figure 5.**
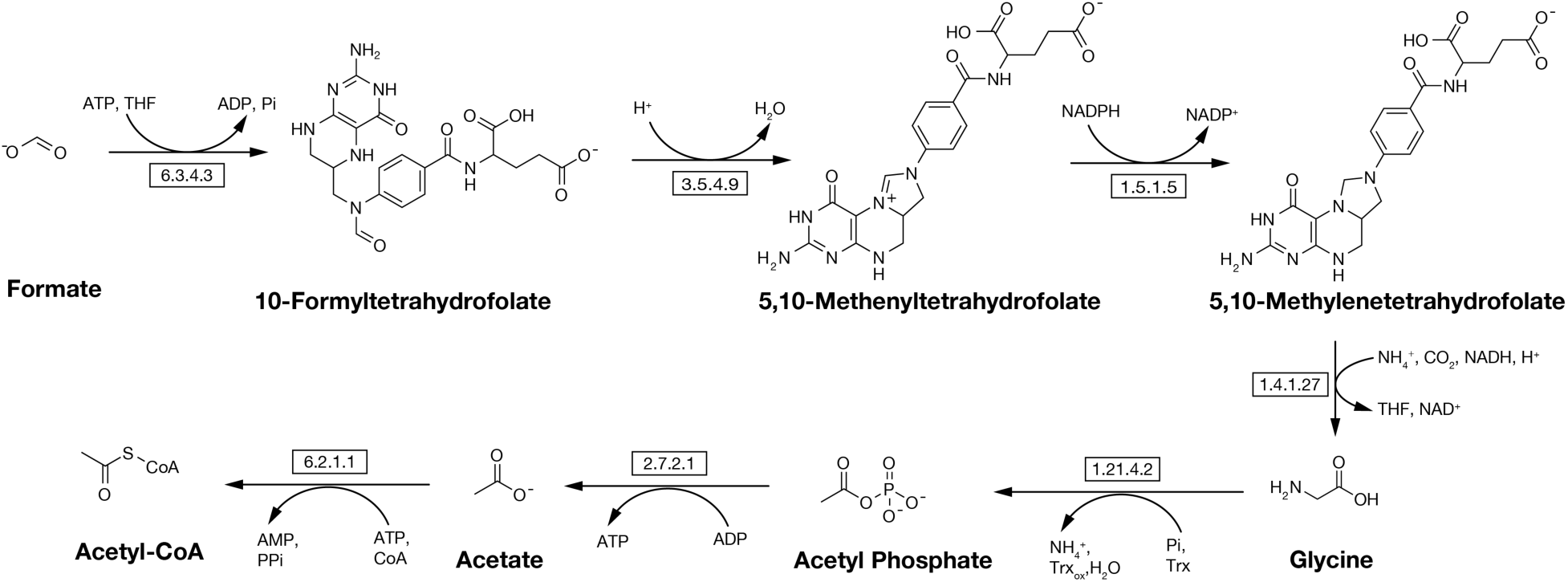
The glycine reductase variant of the reductive glycine pathway (rGly-Gly). The rGly-Gly pathway is a naturally-occurring, selenium-dependent, oxygen-sensitive formate assimilation pathway variant found in microbes known to degrade amino acids and purines^42^. Full details of the reactions shown here can be found in **Dataset S1D**. ATP, NAD(P)H, CO_2_ and HCO_2_^-^ requirements for these pathways are calculated by P4-_R_G_LY_-G_LY_._PY_ in the GitHub repository associated with this article^47^.

**Figure 6.**
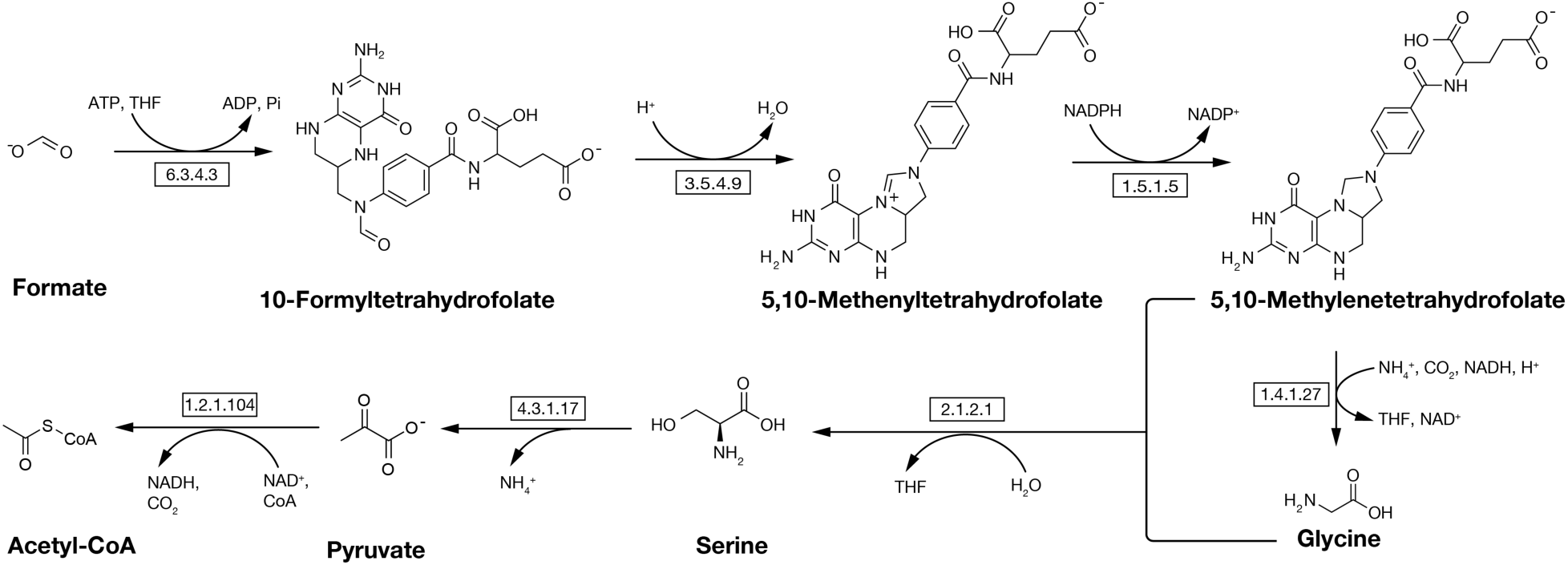
The serine variant of the reductive glycine pathway (rGly-Ser). The rGly-Ser pathway variant is a naturally-occurring, selenium-independent, oxygen-tolerant formate assimilation pathway that produces acetyl-CoA through a serine intermediate^42^. Full details of the reactions shown here can be found in **Dataset S1E**. ATP, NAD(P)H, CO_2_ and HCO_2_^-^ requirements for these pathways are calculated by P5-_R_G_LY_-S_ER_._PY_ in the GitHub repository for this article^47^.

**Figure 7.**
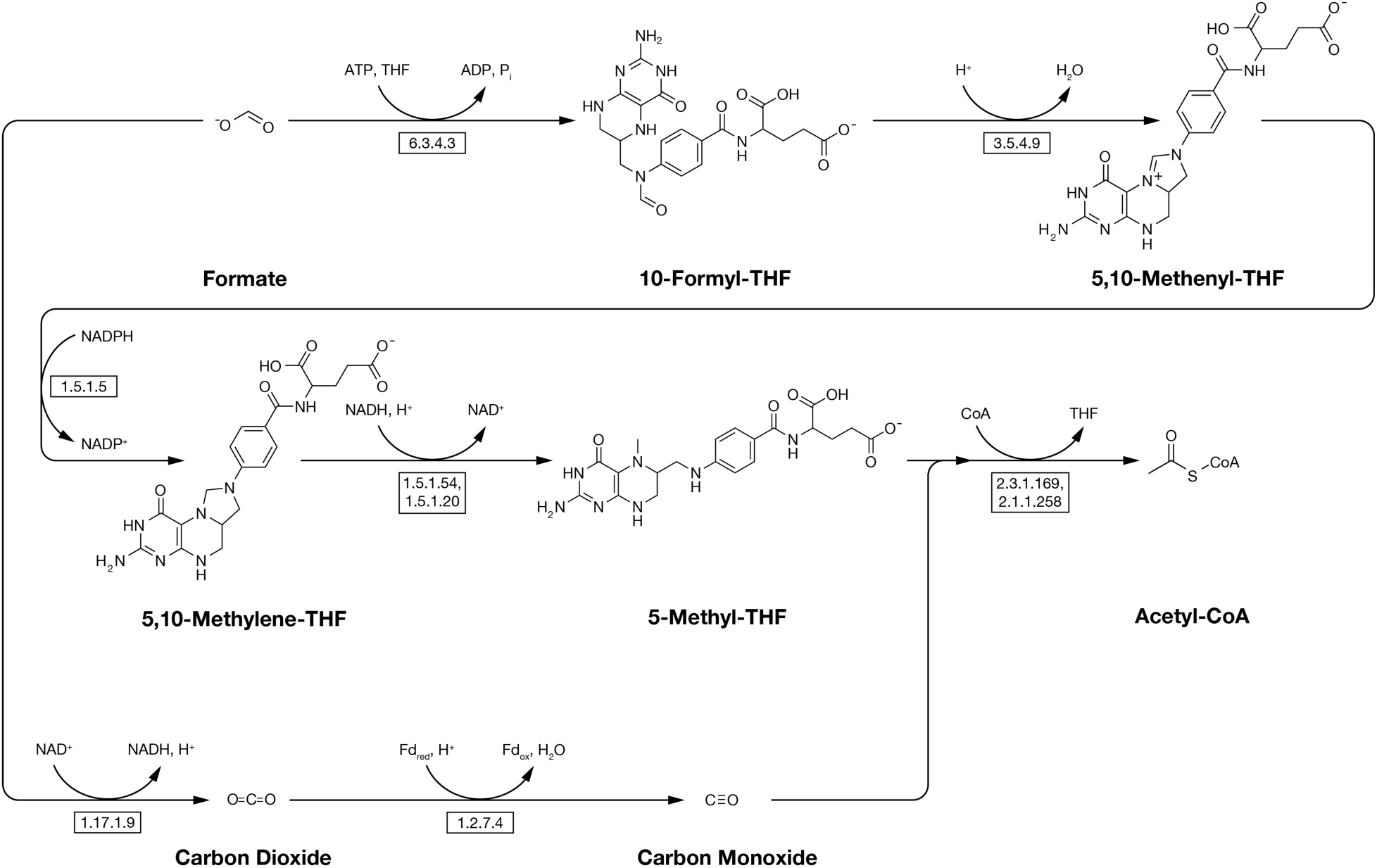
The Wood-Ljungdahl (WL) Pathway. The WL pathway (also known as the reductive acetyl-CoA pathway) is a natural carbon-fixation pathway that functions in anaerobic conditions. The pathway allows for formate assimilation via oxidation of formate to CO_2_ by formate dehydrogenase (E.C. 1.17.1.9). Full details of the reactions shown here can be found in **Dataset S1F**. ATP, NAD(P)H, CO_2,_ and HCO_2_^-^ requirements for these pathways are calculated by P6-FDH-WL._PY_ in the GitHub repository for this article^47^.

Each butanol biosynthesis pathway is comprised of two modules: the first module converts formate (or CO_2_) to acetyl-CoA, and the second module converts acetyl-CoA to butanol. The reactions in the CO_2_-fixation and C_1_- assimilation pathways were used as first modules, and the butanol synthesis pathway (**Figure 8**) was used as the second module in every case.

**Figure 8.**
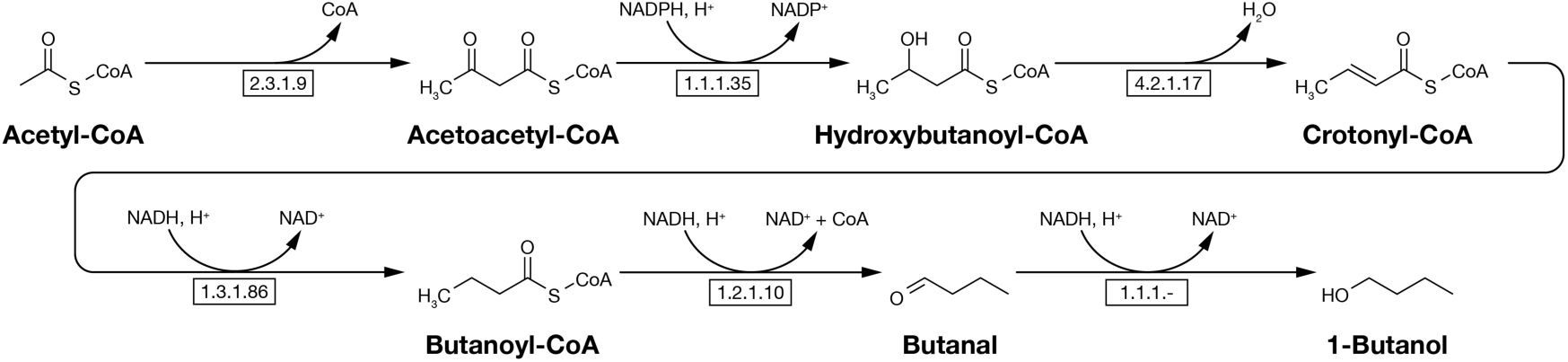
Butanol synthesis pathway. This pathway converts acetyl-CoA produced by CO_2_-fixation or formate-assimilation into 1-butanol. Adapted from Adesina *et al*.^69^. Full details of the reactions shown here can be found in **Dataset S1G**. All computer codes to calculate ATP, NAD(P)H, CO_2,_ and HCO_2_^-^ requirements include this pathway.

ATP, NAD(P)H, formate, and CO_2_ requirements for synthesis of 1-butanol were found by flux balance analysis applied to each two-module pathway analyzed (**Table 2**). Reactions for all pathways were compiled using information from Bar-Even^42^ and data from the Kyoto Encyclopedia of Genes and Genomes (KEGG) database^48–50^. Full reaction lists for each pathway are given in **Datasets S1A** to **S1G**.

We find that formate assimilation can enable energy conversion efficiencies well above the theoretical ceiling of photosynthesis even when formate supplies both the carbon and the electrons. The serine variant of the reductive glycine pathway is close to the best available on efficiency while being the most tractable of the high-efficiency options to engineer.

### The Least Efficient Formate Assimilation Route Converts Sunlight to Butanol at 9.3%, Beating the 8% Upper Limit of Photosynthesis

Formate assimilation routes reach 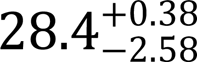 to 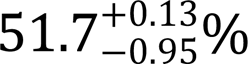 electrical conversion efficiency, corresponding to solar-to-fuel conversion efficiencies of 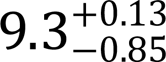 to 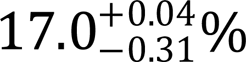. When supplied by a state-of-the-art formate electrolyzer and a photovoltaic operating at the Shockley-Queisser limit of 32.9%^52^, every route in this range exceeds the 8% theoretical ceiling of photosynthesis (**Figure 9A**). The least efficient combination we considered is the glyceraldehyde variant of the formolase pathway supplied with both carbon and electrons by formate (P2a-Form-Glyc), at 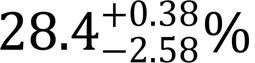 (bar 3 in **Figure 9A**). The most efficient is the Wood-Ljungdahl pathway powered by H_2_ oxidation (P6-FDH-WL), at 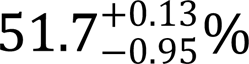 (bar 27).

**Figure 9.**
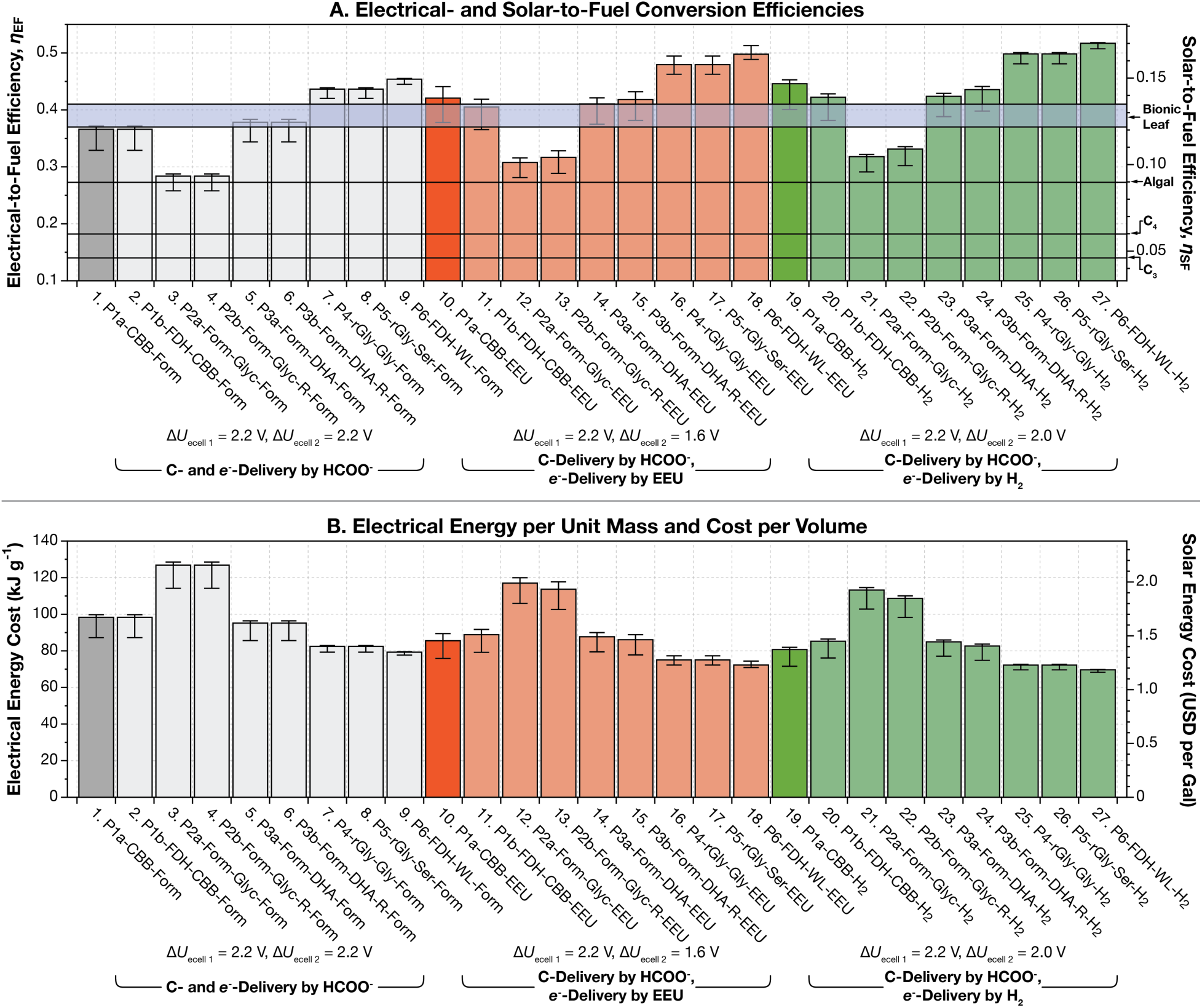
Energy conversion efficiency for carbon assimilation and formate fixation pathways. (**A**) Energy conversion efficiency of butanol production for the Calvin cycle CO_2_-fixation pathway and five formate-assimilation pathways considered in this article. All bars except 1, 10, and 19 show the efficiency of systems supplied with carbon via electrochemically-produced formate. Bars 1, 10, and 19 (in the darkest colors) show the efficiency of systems that fix atmospheric CO_2_ through the Calvin (CBB) cycle. Bars 1 to 9 in grey show the conversion efficiency for systems supplied with electrons through oxidation of formate; bars 10 to 18 in red are results for systems that receive electrons through extracellular electron uptake (EEU); while bars 19 to 27 in green are for systems that receive electrons through H_2_-oxidation. The left axis shows the electrical-to-fuel conversion efficiency. The right axis shows the maximum possible solar-to-fuel conversion efficiency, which assumes that the electricity is supplied by a single-junction, Si photovoltaic operating at the Shockley-Queisser limit of 32.9%^52^. (**B**) Energy costs for butanol production. The left axis shows the electrical energy needed to produce a gram of butanol. The right axis shows the cost to produce one US gallon of butanol, assuming a solar electricity cost of 2¢ per kilowatt hour, the current US Department of Energy SunShot target for utility-scale solar^56^. As described previously^2^, the biggest source of uncertainty in the calculations originates from the value for the potential difference across the cell’s inner membrane, Δ*U*_membrane_. The bars are plotted using the most likely value for the Δ*U*_membrane_, 140 mV (BNIDs 109774, 103386, 109775). The error bars reflect Δ*U*_membrane_ values of 80 mV (BNID 104082) and 270 mV (BNID 107135). This figure can be reproduced with the fig-efficiency-sota.py script in the GitHub repository for this article^47^. State of the art electrolyzer parameters are shown in **Table S2**.

For comparison, the most efficient form of photosynthesis, algal photosynthesis under ideal conditions, has a theoretical upper limit of approximately 8%^53^, and the global average for terrestrial photosynthesis, calculated from satellite measurements of net primary productivity, is closer to 0.1%^11^. The worst combination in our analysis therefore exceeds the theoretical best of the most efficient natural system by roughly 15%, and the best exceeds it by a factor of two.

The comparison here is between two theoretical ceilings, neither of which has been achieved at scale. Plants can approach their own ceilings under favorable conditions: the highest short-term conversion efficiencies measured in C_3_ crops are ≈ 3.5%^54^ against a C_3_ theoretical ceiling of 4.6%^53^, and *Miscanthus* × *giganteus* grown in southern England sustained 3.9% over a full season against a C_4_ theoretical ceiling of about 6%^53,55^. What photosynthesis seems unable to do is sustain that efficiency over large areas, which is why the global average is closer to 0.1%^11^. The bet that EMP proposes is that an engineered system can hold its upper-limit efficiency at scale, because that efficiency depends on the electrolyzer and the engineered organism rather than on soil, water, and weather. If it can, then the land area required to capture a given amount of solar energy falls by a large factor, and a synthetic carbon-fixing system could operate alongside the natural biosphere rather than displacing it.

The same results expressed as energy cost are shown in **Figure 9B**. Producing a gram of butanol requires between 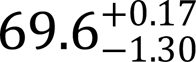 kilojoules (P6-FDH-WL-H_2_, bar 27) and 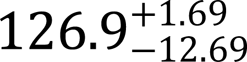 kJ (P2a-Form-Glyc-Form, bar 3). Supplied with solar electricity at 2¢ per kilowatt hour, the US Department of Energy SunShot target for utility-scale solar^56^, this corresponds to 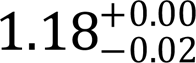 to 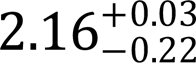 USD per US gallon of butanol.

### Two Sanity Checks Build Confidence in the Calculation

Before comparing pathways, we checked our framework against two cases where the answer was intuited in advance. First, the overall conversion efficiency is identical when formate is used purely as an electron delivery mechanism and when it is used as a carbon and electron delivery mechanism. In our theory, formate supplied for carbon delivery is produced in one electrochemical cell while formate supplied for electron delivery is produced in another (**Figure 1B**), whereas in practice a single cell would serve both. This separation makes formate straightforward to include in the framework but introduces an opportunity for error. To test it, we compared the Calvin cycle fixing atmospheric CO_2_ with formate used purely as an electron mediator (P1a-CBB-Form) against the Calvin cycle supplied with both carbon and electrons by formate, with the carbon released by formate dehydrogenase and refixed by RuBisCO (P1b-FDH-CBB-Form). These should be indistinguishable, and they are: both give 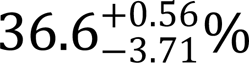 electrical conversion efficiency (bars 1 and 2 in **Figure 9A**). As an aside, while such a distinction seems a little far-fetched (RuBisCO simply refixes the CO_2_ released by oxidation of formate), it is not entirely inconceivable. RuBisCO already discriminates strongly against ^13^C, by 29 to 30 per mil^57^, which is why photosynthetic biomass is isotopically light. One could imagine a variant engineered to be hyper-selective, fixing only ^12^CO_2_ from the atmosphere, supplied with formate made from ^13^CO_2_ so that the carbon released by its oxidation passes through untouched. Formate would then serve purely as a mediator. Whether anyone would wish to construct such a system is another matter.

Second, there is no efficiency gain when formate is used to supply electrons to re-capture carbon lost in butanol synthesis. Both formolase pathways release CO_2_ during the conversion of pyruvate to acetyl-CoA. The glyceraldehyde variant requires eight formate molecules per butanol, losing four carbons for every four incorporated; the dihydroxyacetone variant requires six, losing two (**Table 2**). We therefore added a variant of each pathway in which the lost CO_2_ is recaptured by running formate dehydrogenase in reverse, at the cost of one NADH. Where formate is also the electron source, this should gain nothing, because re-reducing one CO_2_ requires oxidizing another formate, which releases a CO_2_ in its place. Our calculation returns exactly zero gain in both cases, as it should.

However, when electrons arrive more cheaply, recycling does pay off. In an EEU-mediated system, recycling raises the efficiency of the glyceraldehyde variant of the formolase pathway from 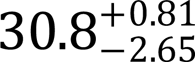 to 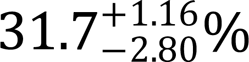 (bars 12 and 13 in **Figure 9A**) and the dihydroxyacetone variant from 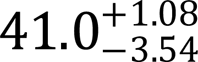 to 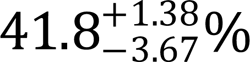 (bars 14 and 15). In an H_2_-mediated system, the efficiency is raised from 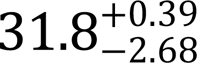 to 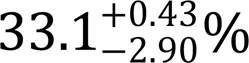 (bars 21 and 22) and 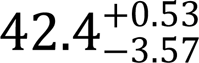 to 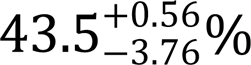 respectively (bars 23 and 24).

The efficiency gains due to carbon recycling in EEU- and H_2_-mediated systems come with a caveat. Running formate dehydrogenase in the reducing direction requires very low formate and very high CO_2_ concentrations, which sacrifices much of what makes formate attractive, including its solubility. We treat these recycling cases as high-efficiency, low-flux scenarios rather than as practical operating points.

### The O_2_-Tolerant rGly-Ser Pathway Is Within 1.9 Points of the O_2_-Sensitive FDH-WL Pathway

The serine variant of the reductive glycine pathway reaches 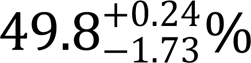 efficiency, within 1.9 points of the best pathway, while being the only high-efficiency option that tolerates oxygen (bars 25 to 27 in **Figure 9A**). The three most efficient pathways we considered are the two variants of the reductive glycine pathway and the Wood-Ljungdahl pathway. In a H_2_-mediated system, the glycine reductase and serine variants of the reductive glycine pathway both reach 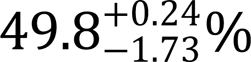 while the Wood-Ljungdahl pathway reaches 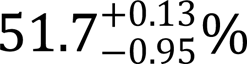 (bars 25 to 27). As seen in our previous works, the ordering is the same in an EEU-mediated system, at 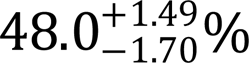 for both reductive glycine variants and 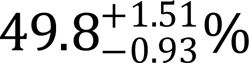 for the Wood-Ljungdahl pathway (bars 16 to 18). In a formate-mediated system, efficiencies are 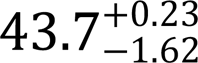 and 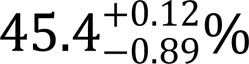 (bars 7 to 9). The gap between the Wood-Ljungdahl pathway and the two reductive glycine variants is only 1.9 points on H_2_, 1.8 on EEU, and 1.7 on formate, in each case smaller than the uncertainty introduced by the membrane potential.

The advantage of the reductive glycine pathway over the Calvin cycle has been demonstrated experimentally. Claassens *et al*. replaced the Calvin cycle in *C. necator* with the reductive glycine pathway and achieved formatotrophic growth^58^, and a subsequent engineered strain reached a biomass yield on formate 17% higher than the wild type using the Calvin cycle, exceeding the yield of any natural formatotroph relying on the Calvin cycle^59^.

The Wood-Ljungdahl and reductive glycine pathways are almost identical in efficiency, but differ substantially in O_2_-tolerance. The Wood-Ljungdahl pathway is highly oxygen-sensitive, which is a serious complication for an EMP organism, because both oxidative phosphorylation and the EEU pathway require oxygen as a terminal electron acceptor. Implementing it would require some form of artificial compartmentalization to shield the pathway from oxygen while still permitting the supply of NAD(P)H and ATP and the export of product. That is a substantial engineering undertaking, and it must be completed before any efficiency measurement can be made at all. Likewise, the glycine reductase variant of the reductive glycine pathway is selenium-dependent and oxygen-sensitive. However, the serine variant is selenium-independent and oxygen-tolerant^42^. This pathway is almost at the top of the efficiency range and can be implemented in an aerobic organism without compartmentalization, making it highly attractive for lab-scale demonstration.

### Formate-Only Systems Lose Three-Quarters of Their Efficiency Between State-of-the-Art and Worst-Case Industrial Electrolyzers, Against Half for H_2_

Having established that the serine variant of the reductive glycine pathway is the most attractive carbon assimilation route, we next ask which electron delivery mechanism to pair with it. **Figure 10** shows this pathway supplied by formate oxidation, EEU, and H_2_-oxidation, at state-of-the-art and at four scaled-up formate production conditions (**Tables S2** and **S3**).

**Figure 10.**
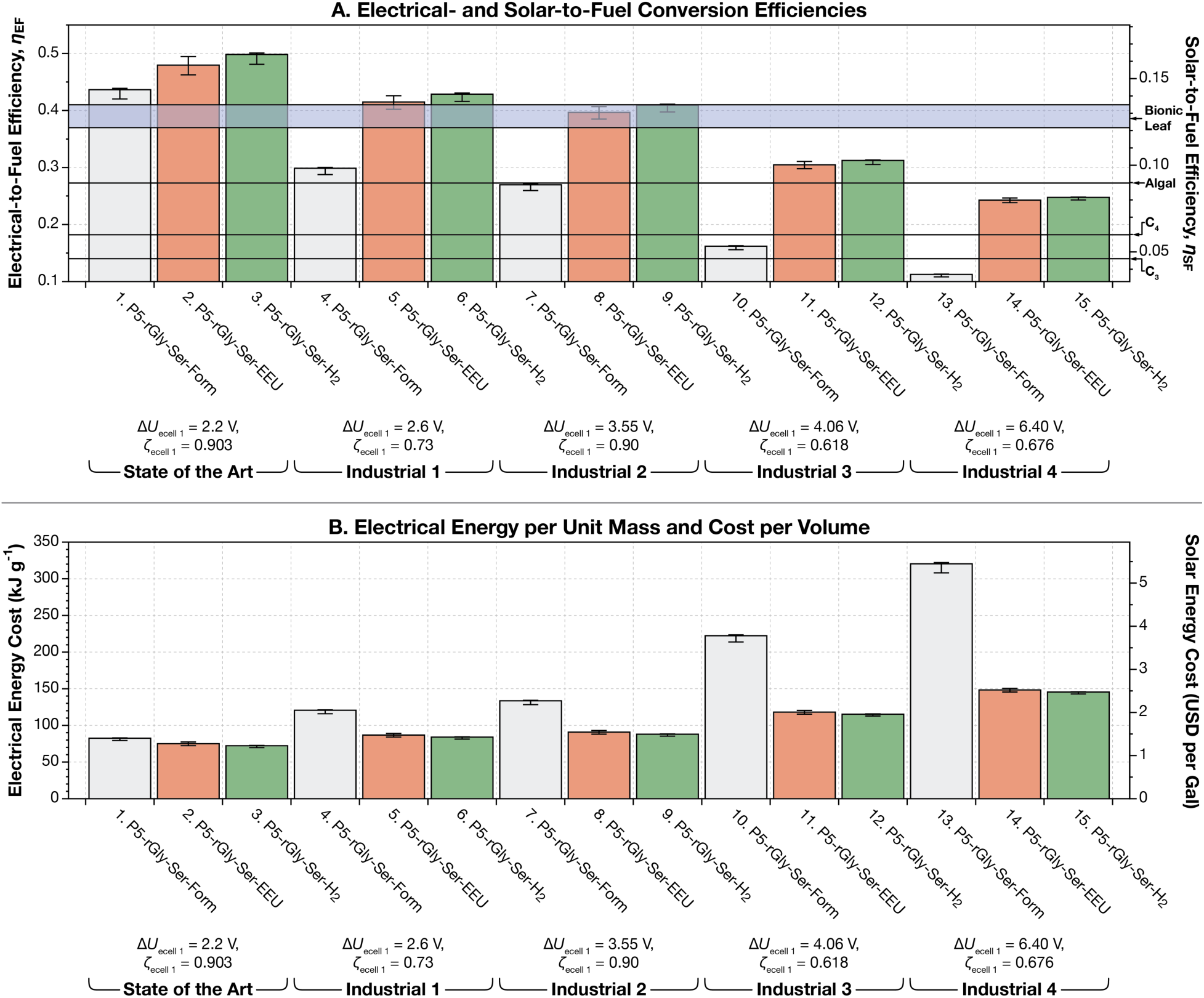
Energy conversion efficiency and energy costs for butanol production with the serine variant of the reductive glycine pathway using state-of-the-art and industrial formate production systems. (**A**) Energy conversion efficiency. The left axis shows the electrical-to-fuel conversion efficiency. The right axis shows the maximum possible solar-to-fuel conversion efficiency, which assumes that the electricity is supplied by a single-junction, Si photovoltaic operating at the Shockley-Queisser limit of 32.9%^52^. (**B**) Energy costs for butanol production. The left axis shows the electrical energy needed to produce a gram of butanol. The right axis shows the cost to produce one US gallon of butanol, assuming a solar electricity cost of 2¢ per kilowatt hour, the current US Department of Energy SunShot target for utility-scale solar^56^. As described previously^2^, the biggest source of uncertainty in the calculations originates from the value for the potential difference across the cell’s inner membrane, Δ*U*_membrane_. The bars are plotted using the most likely value for the Δ*U*_membrane_, 140 mV (BNIDs 109774, 103386, 109775). The error bars reflect Δ*U*_membrane_ values of 80 mV (BNID 104082) and 270 mV (BNID 107135). This figure can be reproduced with the fig-efficiency-ind.py script in the GitHub repository for this article^47^. State of the art and industrial electrolyzer parameters are shown in **Tables S2** and **S3**.

At state-of-the-art conditions the three mechanisms are closer than their reputations suggest. Formate oxidation gives 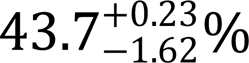 electrical conversion efficiency, EEU 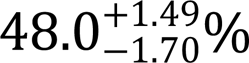, and H_2_-oxidation 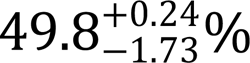 (bars 1 to 3 in **Figure 10A**). The penalty for using formate as both the carbon and the electron source is therefore 6.2 percentage points relative to H_2_, or a relative loss of 12%. Given the difference in experimental tractability between a flask containing formate and a bioelectrochemical reactor requiring a hypoxic chamber and a mediator recycling loop, this seems like a modest price.

The picture changes at scale. Using the whole-cell voltages reported for scaled-up CO_2_-to-formate electrolyzers, efficiency falls in every case, but it falls much faster for the formate-mediated system. At the best scaled-up condition, 2.60 V^60^, formate-mediated efficiency drops to 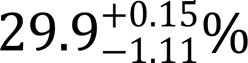 while H_2_ holds at 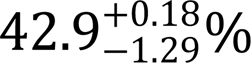 (bars 4 and 6 in **Figure 10A**). At the highest voltage reported, 6.40 V^61^, formate-mediated efficiency falls to 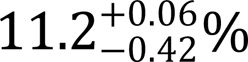 against 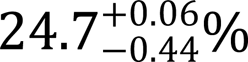 for H_2_- (bars 13 and 15). Across this range the formate-mediated system loses three-quarters of its efficiency while the H_2_-mediated system loses half.

The corresponding energy costs are shown in **Figure 10B**. At state-of-the-art conditions, formate-only operation requires 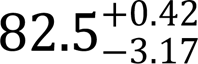 kJ per gram of butanol (bar 1) against 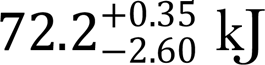 kJ for H_2_ (bar 3), or 1.40 against 1.23 USD per US gallon. At the best scaled-up condition these rise to 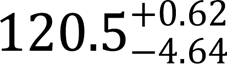 kJ and 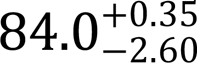 kJ (bars 4 and 6), or 2.05 against 1.43 USD per gallon. At the highest voltage reported, formate-only operation requires 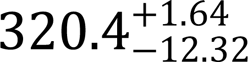 kJ per gram (bar 13), or 5.45 USD per gallon, against 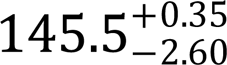 kJ and 2.47 USD for H_2_ (bar 15).

This asymmetry follows directly from the redox limitation described above. Formate supplies two electrons per carbon and butanol requires six, so a system drawing all of its electrons from formate must reduce roughly three times as much CO_2_ at the electrolyzer as one drawing its carbon from formate alone. Every increase in the CO_2_ reduction cell voltage is therefore paid approximately three times over. A system that sources its electrons elsewhere pays that voltage only for the carbon it actually incorporates.

The practical consequence is that the choice of electron delivery mechanism is not fixed by the numbers at state of the art, but becomes decisive at scale. On current laboratory electrochemistry, a formate-only system is a reasonable choice, trading 6.2 points of efficiency for a substantial simplification of the experimental workflow. On the electrochemistry currently available at scale, a formate-only system approaches the theoretical ceiling of photosynthesis rather than exceeding it comfortably, while EEU and H_2_-mediated systems remain well clear.

### Whether the rGly-Ser Carbon Module Can Be Repurposed for a Different Electron Source Is the Central Uncertainty, and Cannot Be Settled by Calculation

The two preceding results pull in opposite directions. Formate-only operation is by far the most tractable route to a working strain, but it is also the route that degrades fastest as electrochemistry moves from the bench to the plant. Given this, is it worth making a high-performing formate-oxidizing, formate-assimilating microbe in the lab?

First, building this organism is no small result. A carbon assimilation pathway demonstrated, characterized, and optimized in a tractable organism stands on its own, whatever is subsequently done with it. Second, the possibility exists that the electron-delivery mechanism could be swapped for H_2_-oxidation or EEU. The assimilation pathway’s demands on the cell are the same in each case, so on paper the two are independent. Whether they are independent in a cell is a different question, and not one this analysis can answer. Whether a strain engineered to assimilate formate using formate-derived electrons can then be converted to use H_2_-oxidation or EEU is an experimental question, and we expect it can only be settled by attempting it.

The bet is therefore worth stating explicitly. If a formate-only strain is built and later proves impossible to convert to EEU or H_2_ electron delivery, is the work useful, or is the strain a dead end? We think it is still useful. While H_2_- and EEU-mediated systems have higher theoretical ceilings, state-of-the-art whole-cell voltages for CO_2_ reduction to formate have fallen from around 3.5 V to 2.2 V in a decade (**Table S2**), and the best scaled-up system now reported, at 2.60 V^60^, is already better than the laboratory state of the art of a few years ago. The scenarios in which formate-only operation performs poorly are those built on the highest-voltage prototypes currently reported, and there is no indication that the improvement in this technology will stop any time soon.

### The Calculation Points Toward a Chassis That Is Aerobic, Fast, and Genetically Tractable

The pathway we identify imposes requirements on the organism that implements it. Because the serine variant of the reductive glycine pathway is oxygen-tolerant, it does not require an anaerobic chassis, and because regenerating ATP by oxidative phosphorylation requires oxygen in any case, an aerobic organism is preferable. The pathway is also short, and it has now been installed in a range of hosts, including *E. coli*^62,63^, *C. necator*^58^, *Saccharomyces cerevisiae*^64^, and *Pseudomonas putida*^65^, so the choice of chassis is not strongly constrained by the pathway itself.

Beyond these, the argument of this article favors an organism in which the design-build-test cycle is fast. If the gap between an upper-limit efficiency and a realized one is closed by iteration, then the rate at which iterations can be performed is as important as the ceiling itself. The recent history of synthetic formatotrophy bears this out. Engineered formatotrophs had doubling times above six hours until the fastest reported strain reached 4.4 hours, close to the roughly four hours of the fastest natural formatotrophs, and a laboratory-evolved *C. necator* running its native Calvin cycle has reached 3.3 hours^66^.

Taken together, these criteria narrow the field considerably. An organism is wanted that is aerobic, grows rapidly, is highly amenable to genetic manipulation, and possesses a native route for taking up electrons, whether by H_2_- oxidation or by EEU. The last of these is the most restrictive, since the organisms in which electron uptake is best characterized are generally not those that grow fastest or engineer most easily.

Several organisms meet part of this description. *C. necator* oxidizes H_2_ natively, is the chassis behind existing commercial EMP systems, and has already been engineered to run the reductive glycine pathway in place of its native Calvin cycle^58^, though it grows more slowly and is less tractable than the standard laboratory hosts. *P. putida* is robust and reasonably tractable, and has been engineered for mediator-based electroactivity through heterologous phenazine biosynthesis^67^, but as with *V. natriegens* this was demonstrated as electron export to an anode, and it has no native uptake capability. *V. natriegens* is a facultative anaerobe, among the fastest-growing microorganisms known, highly amenable to genetic manipulation, and possesses a characterized Mtr-type extracellular electron transfer conduit^32,33^.

## Conclusion

We set out to determine which combination of carbon assimilation pathway and electron delivery mechanism is worth building. The calculation gives a clear answer on the first and a conditional one on the second.

On the pathway, the serine variant of the reductive glycine pathway is the strongest candidate. It reaches 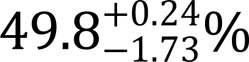 electrical conversion efficiency on H_2_-oxidation, within 1.9 points of the Wood-Ljungdahl pathway, a gap smaller than the uncertainty in our own calculation. Unlike the Wood-Ljungdahl pathway and the glycine reductase variant, it is oxygen-tolerant, so it can be implemented in an aerobic organism without compartmentalization. It is also selenium-independent, short, and has already been installed in several hosts.

On the electron delivery mechanism, the answer depends on the electrochemistry available. At state-of-the-art conditions, drawing both carbon and electrons from formate costs 6.2 percentage points against H_2_-oxidation, which is a modest price for a workflow that requires no electrode, no gas handling, and no hypoxic chamber. At the highest whole-cell voltages currently reported for scaled-up CO_2_-to-formate electrolyzers, the same system falls to 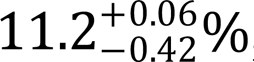, below the theoretical ceiling of photosynthesis, while H_2_-mediated systems remain well clear.

This makes the recommendation conditional in a way that can be tested. A formate-only system is the right first target if CO_2_-to-formate electrochemistry continues to improve, and the wrong one if it does not. The trajectory favors the first case: whole-cell voltages have fallen from around 3.5 V to 2.2 V in a decade, and the best scaled-up system now reported operates at 2.60 V, better than the laboratory state of the art of a few years ago. If that trend halts, the argument for building a formate-only system first weakens considerably, and the case for paying the additional engineering cost of extracellular electron uptake or H_2_-oxidation at the outset becomes correspondingly stronger.

Two questions remain open, and neither can be settled by calculation. The first is whether a carbon assimilation pathway engineered to run on formate-derived electrons can subsequently be converted to another electron source, or whether the two are entangled in ways this analysis cannot see. The second is how much of the gap between an upper-limit efficiency and a realized one can be closed by iteration, and how quickly. Both are experimental questions, and both argue for beginning in an organism where iteration is fast.

## End Notes

### Code Availability

All code used in calculations in this article is available at https://github.com/barstowlab/article-033-formate-efficiency^47^.

### Materials & Correspondence

Correspondence and material requests should be addressed to B.B..

### Author Contributions

Conceptualization, D.S., D.A.S, and B.B.; Methodology, D.S., R.A., F.S., T.J.S. D.A.S., and B.B.; Investigation, D.S., R.A., L.O.A., S.S., and B.B.; Writing—Original draft, D.S., R.A., L.O.A., and B.B.; Writing—Review and editing, D.S., R.A., L.O.A., and B.B.; Funding acquisition, D.S., R.A., L.O.A., and B.B.; Resources, B.B.; Supervision, B.B. and D.A.S.; Data curation, B.B.; Visualization, D.S., R.A., L.O.A., and B.B.; Formal analysis, D.S., R.A., L.O.A., and B.B..

## Acknowledgments

This work was supported by Cornell University startup funds, a Career Award at the Scientific Interface from the Burroughs-Welcome Fund, a gift from Mary Fernando Conrad and Tony Conrad to B.B.. D.A.S. was supported by a Cornell Energy Systems Institute fellowship. D.S. was supported by the Cornell Engineering Learning Initiative. R.A. was supported by a fellowship from Saudi Aramco. L.O.A. was supported by a fellowship from the UK Biochemical Society.

