## Supplemental Information for "Thermodynamic, Electrochemical and Practical Constraints on Electromicrobial Formate Assimilation"

### Supplementary Information Tables

**Table S1.** Symbols used in this article.

**Table S2.** Electrochemical parameters for state-of-the-art laboratory scale $CO_2$ to formate/formic acid reduction.

**Table S3.** Electrochemical parameters for scaled-up $CO_2$ to formate/formic acid reduction.

**Table S4.** Electrochemical parameters for state-of-the-art laboratory scale $H_2$ production cells.

**Table S5.** Electrochemical parameters for scaled-up $H_2$ production cells.

### Supplementary Information Datasets

**Dataset S1.** Enzymatic reactions for conversion of $CO_2$ and formate/formic acid to butanol.

| **Symbol** | **Unit** | **Description** |
|---|---|---|
| $E_p$ | J molecule$^{-1}$ | Energy carried per product (*e.g*, fuel) molecule. |
| $\dot{N}_p$ | molecule s$^{-1}$ | Product (*e.g.*, fuel) molecules produced per second by electromicrobial production system. |
| $N_A$ | molecule Mol$^{-1}$ | Avogadro constant. |
| F | A s Mol$^{-1}$ | Faraday Constant. |
| $P_{e,\ total}$ | J s$^{-1}$ | Total electrical power input into electromicrobial production system. |
| $M_p$ | g Mol $^{-1}$ | Molecular weight of the fuel molecule. |
| $e$ | A s | Fundamental charge. |
| $\nu_{ep}$ | # | Number of electrons needed for synthesis of a product (*e.g*, fuel) molecule. |
| $\Delta U_{ecell\ 1}$ | V | Potential difference across $CO_2$-reduction electrochemical cell (cell 1). |
| $\Delta U_{ecell\ 2}$ | V | Potential difference across bio-electrochemical cell (cell 2). |
| $I_{ecell\ 1}$ | A | Current through $CO_2$-reduction electrochemical cell (cell 1). |
| $I_{ecell\ 2}$ | A | Current through bio-electrochemical cell (cell 2). |
| $j_{ecell\ 1}$ | mA cm$^{-2}$ | Current density through $CO_2$-reduction electrochemical cell (cell 1). |
| $\nu_{e,\ add}$ | # | Number of electrons needed to convert a $CO_2$-reduction product (*e.g*, formate or other $C_1$ or $C_2$ compound) to a molecule of final product. |
| $\nu_r$ | # | Number of primary reduction products to make a molecule of final product. |
| $\nu_{er}$ | # | Number of electrons to reduce $CO_2$ to a primary reduction product. |
| $\nu_{Cr}$ | # | Number of carbon atoms per primary reduction product. |
| $\zeta_{I2}$ | # | Faradaic efficiency of the bio-electrochemical cell. |
| $\zeta_{I1}$ | # | Faradaic efficiency of the primary abiotic cell. |
| $\zeta_C$ | # | Carbon transfer efficiency from cell 1 to cell 2. |
| $\nu_{p,\ NADH}$ | # | Number of NAD(P)H molecules needed to make a final product molecule. |
| $\nu_{p,\ Fd}$ | # | Number of Ferredoxin molecules needed to make a final product molecule. |
| $\nu_{p,\ ATP}$ | # | Number of ATP molecules needed to make a final product molecule. |
| $\Delta G_{ATP/ADP}$ | J | Free energy for regeneration of ATP. |
| $\Delta U_{membrane}$ | V | Inner membrane potential difference. |
| $U_{H2}$ | V | Standard potential of proton reduction to $H_2$. |
| $U_{acceptor}$ | V | Standard potential of terminal electron acceptor reduction. |
| $U_Q$ | V | Redox potential of the inner membrane electron carrier. |
| $U_{NADH}$ | V | Standard potential of NADH. |
| $U_{Fd}$ | V | Standard potential of Ferredoxin. |
| $C_{EF}$ | J g$^{-1}$ | Electrical energy cost per unit mass. |
| $C_{SF}$ | ¢ g$^{-1}$ | Financial cost efficiency per unit mass. |
| $\eta_{EF}$ | % | Electrical to product (*e.g*, fuel) energy conversion efficiency. |
| $\eta_{SF}$ | % | Solar to product (*e.g*, fuel) energy conversion efficiency. |

**Table S1.** Symbols used in this article.

| Description | Cell Voltage, $\Delta U_{ecell}$ (V) | Current Density, $j_{ecell}$ (mA cm$^{-2}$) | Faradaic Efficiency, $\zeta$ (%) | Temperature (°C) | Stability (hours) | References |
|---|---|---|---|---|---|---|
| Cu@BOC catalyst (Cu-stabilised $Bi_2O_2CO_3$) in 100 cm$^2$ alkaline Membrane Electrode Assembly (MEA). | 2.2 | 100 | 90.3 | RT | NR | [ZhangH2025a] |
| Cu@BOC – as above. | 2.3 | 200 | 93.4 | RT | NR | [ZhangH2025a] |
| Cu@BOC – as above. | 2.4 | 300 | 92.5 | RT | NR | [ZhangH2025a] |
| Cu@BOC – as above. | 2.5 | 400 | 90.2 | RT | NR | [ZhangH2025a] |
| Pitted bismuth nanosheet catalyst in a zero-gap MEA, 1 M KOH electrolyte, humidified $CO_2$ to cathode. | 2.84 | 200 | > 90 | NR | 110 | [Yuan2022a] |
| Core-shell Cu/Bi nanowires on 3D Cu foam in MEA with cation-exchange membrane (CEM). | 2.87 | 200 | 90 | NR | 8000 | [LiJJ2026a] |
| Bi/C nanoparticle catalyst on Gas Diffusion Electrode (GDE) in a gas-phase filter-press MEA. | 3.02 | 45 | 90 | 25 ± 1 | NR | [Diaz-Sainz2020a] |
| Turing-structured $Sb_{0.1}Sn_{0.9}O_2$ catalyst in a commercial 5 cm$^2$ zero-gap MEA (Dioxide Materials). | 3.3 | 500 | 82 | NR | 200 | [Ye2025a] |
| Bi/C catalyst on GDE in gas-phase filter press, Sustainion Anion-Exchange Membrane (AEM), 1 M KOH, $IrO_2$-MMO (Mixed Metal Oxide) anode. | 3.4 | 200 | 93 | RT | NR | [Diaz-Sainz2021a] |
| Vertical Bi nanosheet catalysts in slim continuous-flow cell, 0.05 M $H_2SO_4$ + 0.5 M $K_2SO_4$ (pH 2); product is formic acid. | 3.44 | 471 | 96.3 | 25 | 50 | [Chi2023a] |
| Sn nanoparticle catalyst on GDE in three-compartment cell (AEM, ion-exchange resin, CEM). | 3.5 | 140 | 94 | RT | 500 | [Yang2017a] |
| Sn plate catalyst in high-pressure bipolar-membrane cell, 40–50 bar $CO_2$; product is a formic acid/formate mixture. | 3.5 | 30 | 90 | 22 ± 1 | NR | [Ramdin2019a] |
| $Bi_2O_3$ catalyst on GDE in three-compartment cell (AEM, ion-exchange resin, CEM). | 3.52 | 200 | 75.9 | RT | 1000 | [YangH2020a] |
| $BiSn_xO_y$ Sn@Bi catalyst in an MEA. | 3.6 | 270 | 94.3 | RT | 160 | [Cheng2025a] |
| $Cu_6Sn_5$ catalyst in a cation-free solid-state-electrolyte MEA, pH 1; product is formic acid. | 3.7 | 100 | 75.9 | RT | 130 | [Yu2024a] |
| Indium mesh cathode in a closed-loop flow-cell stack of three 109 cm$^2$ cells in series, 1 M KOH anolyte with a Nafion 324 membrane, powered directly by a silicon photovoltaic array under natural AM1.5 sunlight; voltage is per cell and is the lowest of the anolytes studied. | 3.7 | 73 – 92 | 67 | NR | NR | [White2014a] |
| Sn plate catalyst in high-pressure bipolar-membrane cell, 40–50 bar $CO_2$; product is a formic acid/formate mixture; as above but at higher cell potential. | 4 | 100 | 65 | 22 ± 1 | NR | [Ramdin2019a] |

**Table S2.** Electrochemical parameters for state-of-the-art laboratory-scale $CO_2$-to-formate/formic acid electrolyzers coupled to an oxygen evolution reaction (OER) from water splitting at the anode. **Abbreviations:** AEM – Anion-Exchange Membrane; CEM – Cation-Exchange Membrane; GDE – Gas Diffusion Electrode; MEA – Membrane Electrode Assembly; MMO – Mixed Metal Oxide; NR – Not Reported. **Note:** Zhang *et al.* [ZhangH2025a] report 130 hours of stable operation, but under methanol oxidation (MOR) at the anode rather than OER; no durability is reported for the $CO_2$ reduction, OER configuration tabulated here.

| **Description** | **Cell Voltage, $\Delta U_{ecell}$ (V)** | **Current Density, $j_{ecell}$ (mA cm$^{-2}$)** | **Faradaic Efficiency, $\zeta$ (%)** | **Stability (hours)** | **References** |
|---|---|---|---|---|---|
| VITO CORE® Sn catalyst on single Gas Diffusion Electrode (GDE), 400 cm$^2$ active area, alkaline flow cell. | 2.60 | 100 | 73 | NR | [Izadi2026a] |
| 3D-Cu/Bi core–shell nanowire catalyst on Cu foam in scaled-up Membrane Electrode Assembly (MEA), 100 cm$^2$ active area, dry $CO_2$ feed, $IrO_2$ anode. | 3.55 | 200 | 90 | 2000 | [LiJJ2026a] |
| Electro Carbon Inc., Sn catalyst on GDE, 1526 cm$^2$ active area per cell, three-compartment flow cells, stack of two cells in series, 4 M KOH catholyte; product is potassium formate. | 4.06 | 200 | 61.8 | NR | [Fink2024a] |
| University of Cantabria + Apria Systems Zero-gap gas-phase prototype, Bi-based catalyst on GDE, 100 cm$^2$ active area, with serpentine flow field. | 6.40 | 200 | 67.6 | NR | [Abarca2025a] |

**Table S3.** Electrochemical parameters for scaled-up $CO_2$-to-formate/formic acid electrolyzers coupled to an oxygen evolution reaction (OER) from water splitting at the anode. **Abbreviations:** GDE – Gas Diffusion Electrode; MEA – Membrane Electrode Assembly; NR – Not Reported. **Note 1:** We are reporting the whole-cell voltage achieved by Li *et al*. [LiJJ2026a] with a non-optimized anode. **Note 2:** Fink *et al*. [Fink2024a] used a dual cell system, but we are reporting mean per-cell voltage. **Note 3:** Operating temperatures are not reported for any entry except Fink *et al.* [Fink2024a], and the remainder are assumed to operate at ambient temperature. Fink *et al.* [Fink2024a] held each electrolyte at 20 to 25 ˚C by continuous cooling with a chiller of up to 3.3 kW capacity.

| Description | Cell Voltage, $\Delta U_{ecell}$ (V) | Current Density $j_{ecell}$ (mA/cm$^2$) | Faradaic Efficiency, $\xi$ (%) | Temperature (°C) | Stability (hours) | References |
|---|---|---|---|---|---|---|
| Capillary-fed electrolysis cell, 27 wt% aqueous KOH drawn through a polyether sulfone separator by capillary action, polytetrafluoroethylene (PTFE) treated anode. | 1.51 | 500 | 100 | 85 | NR | [Hodges2022a] |
| Anion Exchange Membrane Water Electrolysis (AEMWE) with gas-permeable QC6$_{50}$BA-2.1 anion-exchange ionomer and $Ni_{0.8}Co_{0.2}O$ anode; voltage is IR-included. Stability at 1,000 mA cm$^{-2}$. | 1.69 | 2,000 | 100 | 80 | 1000 | [Liu2024a] |
| AEMWE five-cell stack, 64 cm² per cell, NiCoO-NiCo/C cathode and CuCoO anode, 1 M KOH; voltage is per cell, 9.25 V across the stack. | 1.85 | 740 | 100 | 50 | 150 | [Park2021a] |
| Bionic Leaf biocompatible water-splitting catalyst pair: cobalt–phosphorus (Co-P) alloy cathode and cobalt phosphate (CoPi) anode on high-surface-area carbon cloth, 4 cm² geometric area per electrode, 36 mM phosphate-buffered minimal medium at pH 7, two-electrode configuration, operated in the presence of *Ralstonia eutropha* (also known as *Cupriavidus necator*). Used in all of our previous theoretical work [Salimijazi2020b, Wise2022a, Marecos2022b, Sheppard2023b, Sheppard2024a]. | 2.0 | NR | 99 | 30 | 384 | [Liu2016a] |
| Ultra-high rate AEMWE with PTP Anion-Exchange Membrane (AEM), PBP ionomer, 3D AEI-CEI-NiFe/Ni-foam anode, Pt/C cathode, 1 M KOH. | 2.30 | 10,000 | 100 | 80 | > 800 | [Zheng2025a] |
| Earlier version of Bionic Leaf: cobalt phosphate (CoPi) anode with an electrodeposited nickel–molybdenum–zinc (NiMoZn) cathode on stainless steel mesh, chloride-free minimal medium at pH 7, two-chamber H cell, operated in the presence of *R. eutropha* (aka *C. necator*); sustained microbial growth required at least 2.7 V. | 2.5 | 4 | NR | RT | 120 | [Torella2015a] |

**Table S4.** Electrochemical parameters for lab-scale and state-of-the-art $H_2$ production cells coupled to an oxygen evolution reaction (OER) from water splitting at the anode. **Abbreviations:** AEM – Anion Exchange Membrane; AEMWE – Anion Exchange Membrane Water Electrolysis; NR – Not Reported; RT – Room Temperature. **Note:** Almost all electrolyzers operate at high temperature with the exception of the Bionic Leaf systems.

| **Description** | **Cell Voltage, $\Delta U_{ecell}$ (V)** | **Current Density $j_{ecell}$ (mA/cm²)** | **Specific Energy Consumption, $E_s$ (kWh kg⁻¹)** | **Temperature (°C)** | **References** |
|---|---|---|---|---|---|
| Bloom Energy Solid Oxide Electrolysis Cell (SOEC), 100 kW prototype system independently tested at Idaho National Laboratory; direct-current stack consumption, achieved with an external steam supply. | 1.38† | NR | 36.7 | NR | [Casteel2025a] |
| Bloom Energy Solid Oxide Electrolysis Cell (SOEC) at commercial scale. | 1.41† | NR | 37.5 | NR | [Bloom2026a] |
| Solid Oxide Electrolysis Cell (SOEC) systems from Sunfire, Bloom Energy and Ceres; alternating-current system consumption at 85 to 88% efficiency on a lower-heating-value basis, conditional on an available steam supply. | 1.39 – 1.47† | NR | 37 – 39 | NR | [Riegraf2025a] |
| Hysata capillary-fed alkaline electrolyser at pre-commercial scale, reported as 98% cell energy efficiency on a higher-heating-value basis. | 1.56† | NR | 41.5 | NR | [Hysata2022a] |
| Commercial Alkaline Water Electrolysis (AWE) and Proton Exchange Membrane (PEM) cells, benchmark polarisation curves reproduced by [Hodges2022a]; 1.77 V corresponds to 500 mA/cm² for the alkaline cell and 1800 mA/cm² for the PEM cell. | 1.77 | 500; 1,800 | 47.1† | 90 | [Debe2012a] |
| Commercialised monopolar Alkaline Water Electrolysis (AWE) units, 25 to 35 wt% KOH, activated Ni-plated or Ni-coated steel cathodes on mild steel or steel anodes, up to 200 kW and 42 m³ $H_2$ per hour. | 1.75 – 1.90 | 134 – 250 | 46.5 – 50.5† | 70 – 82 | [Zeng2010a] |
| Commercialised Proton Exchange Membrane (PEM) electrolyser units, 7 to 51 cells per stack, 1 kA maximum. | 1.7 – 2.0 | 500 – 1,075 | 45.2 – 53.2† | 65 – 80 | [Zeng2010a] |
| Alkaline Water Electrolysis (AWE) systems at megawatt to gigawatt scale; alternating-current system consumption. | 1.84 – 1.88† | NR | 49 – 50 | NR | [Riegraf2025a] |
| Proton Exchange Membrane Water Electrolysis (PEMWE), typical range of commercial specifications. | 1.8 – 2.2 | 600 – 2,000 | 47.9 – 58.5† | 50 – 80 | [Smolinka2011a, Carmo2013a] |
| Proton Exchange Membrane Water Electrolysis (PEMWE) systems at megawatt to gigawatt scale; alternating-current system consumption. | 1.96 – 2.07† | NR | 52 – 55 | NR | [Riegraf2025a] |
| Alkaline Water Electrolysis (AWE), typical range of commercial specifications. | 1.8 – 2.4 | 200 – 400 | 47.9 – 63.8† | 60 – 80 | [Smolinka2011a, Carmo2013a] |
| Siemens Energy Silyzer 200 Proton Exchange Membrane (PEM) electrolyser, 5 MW, 20 kg $H_2$ per hour, the largest operating PEM electrolyser system at the time of publication; alternating-current system consumption. | 2.26† | NR | 60 | NR | [Siemens2019a] |

**Table S5.** Electrochemical parameters for scaled-up $H_2$ production cells coupled to an oxygen evolution reaction (OER) from water splitting at the anode. **Abbreviations:** AWE – Alkaline Water Electrolysis; NR – Not Reported; PEM – Proton Exchange Membrane; PEMWE – Proton Exchange Membrane Water Electrolysis; SOEC – Solid Oxide Electrolysis Cell. **Note 1:** Values marked † are derived rather than reported. Specific energy consumption and cell voltage are interconvertible assuming unit faradaic efficiency. One kilogram of $H_2$ is equivalent to $9.572 \times 10^7$ coulombs, and energy is equivalent to charge × voltage, meaning that $\Delta U_{ecell} = 0.0376 \times E_s$.

### Supplementary Information References


[Abarca2025a] J.A. Abarca, C. Gonzalez-Fernandez, C.E. Peralta, A. Arruti, E. Santos, G. Diaz-Sainz, and A. Irabien, "Prototype Validation of a Large-Scale $CO_2$-to-Formate Zero-Gap Electrolyzer", *ChemSusChem* 18, e202501116 (2025). doi:10.1002/cssc.202501116.

[Adesina2017a] O. Adesina, I.A. Anzai, J.L. Avalos, and B. Barstow, "Embracing Biological Solutions to the Sustainable Energy Challenge", *Chem* 2, 20–51 (2017). doi:10.1016/j.chempr.2016.12.009.

[Bar-Even2016a] A. Bar-Even, "Formate Assimilation: The Metabolic Architecture of Natural and Synthetic Pathways", *Biochemistry* 55, 3851–3863 (2016). doi:10.1021/acs.biochem.6b00495.

[Barstow2026a] B. Barstow, "Formate Efficiency 1 Release for Zenodo" (2026). doi:10.5281/zenodo.22903406.

[Bloom2026a] Bloom Energy, "An Efficient Electrolyzer for Clean Hydrogen", Bloom Energy. https://www.bloomenergy.com/bloomelectrolyzer/.

[Carmo2013a] M. Carmo, D.L. Fritz, J. Mergel, and D. Stolten, "A comprehensive review on PEM water electrolysis", *International Journal of Hydrogen Energy* 38, 4901–4934 (2013). doi:10.1016/j.ijhydene.2013.01.151.

[Casteel2025a] M. Casteel, T.L. Westover, A. Shigrekar, T. Olowu, A. Ta, A. Lavernia, A. Zargari, and B. Cheldelin, "Ultra-high efficiency hydrogen production using a large-scale solid oxide electrolysis cell system", *International Journal of Hydrogen Energy* 157 (2025). doi:10.1016/j.ijhydene.2025.150283.

[Cheng2025a] Z. Cheng, J. Song, L. Liu, C. Qiu, L. Wang, and J. Wang, "Amorphous $BiSn_xO_y$ for Efficient $CO_2$ Electroreduction to Formate via *In Situ* Doping", *Advanced Science* 13, e22395 (2026). doi:10.1002/advs.202522395.

[Chi2023a] L.P. Chi, Z.Z. Niu, Y.C. Zhang, X.L. Zhang, J. Liao, Z.Z. Wu, P.C. Yu, M.H. Fan, K.B. Tang, and M.R. Gao, "Efficient and stable acidic $CO_2$ electrolysis to formic acid by a reservoir structure design", *Proceedings of the National Academy of Sciences* 120, e2312876120 (2023). doi:10.1073/pnas.2312876120.

[Debe2012a] M.K. Debe, S.M. Hendricks, G.D. Vernstrom, M. Meyers, M. Brostrom, M. Stephens, Q. Chan, J. Willey, M. Hamden, C.K. Mittelsteadt, C.B. Capuano, K.E. Ayers, and E.B. Anderson, "Initial Performance and Durability of Ultra-Low Loaded NSTF Electrodes for PEM Electrolyzers", *Journal of The Electrochemical Society* 159, K165–K176 (2012). doi:10.1149/2.065206jes.

[Diaz-Sainz2020a] G. Díaz-Sainz, M. Alvarez-Guerra, B. Ávila-Bolívar, J. Solla-Gullón, V. Montiel, and A. Irabien, "Improving trade-offs in the figures of merit of gas-phase single-pass continuous $CO_2$ electrocatalytic reduction to formate", *Chemical Engineering Journal* 405 (2021). doi:10.1016/j.cej.2020.126965.

[Diaz-Sainz2021a] G. Díaz-Sainz, M. Alvarez-Guerra, and A. Irabien, "Continuous electroreduction of $CO_2$ towards formate in gas-phase operation at high current densities with an anion exchange membrane", *Journal of CO2 Utilization* 56 (2022). doi:10.1016/j.jcou.2021.101822.

[Fink2024a] A.G. Fink, F. Navarro-Pardo, J.R. Tavares, and U. Legrand, "Scale-up of electrochemical flow cell towards industrial $CO_2$ reduction to potassium formate", *ChemCatChem* 16 (2024). doi:10.1002/cctc.202300977.

[Hodges2022a] A. Hodges, A.L. Hoang, G. Tsekouras, K. Wagner, C.Y. Lee, G.F. Swiegers, and G.G. Wallace, "A high-performance capillary-fed electrolysis cell promises more cost-competitive renewable hydrogen", *Nature Communications* 13, 1304 (2022). doi:10.1038/s41467-022-28953-x.

[Hysata2022a] Hysata, "Hysata's electrolyser breaks efficiency records", Hysata (2022). https://hysata.com/news/hysatas-electrolyser-breaks-efficiency-records-enabling-world-beating-green-hydrogen-cost/.

[Izadi2026a] P. Izadi, S. Varhade, C. Schneider, P. Haus, C. Singh, A. Guruji, D. Pant, and F. Harnisch, "Scaling up electrochemical $CO_2$ reduction to formate through comparative reactor analysis", *Industrial Chemistry & Materials* 4, 260–275 (2026). doi:10.1039/d5im00056d.

[Kanehisa2000a] M. Kanehisa and S. Goto, "KEGG: Kyoto Encyclopedia of Genes and Genomes", *Nucleic Acids Research* 28, 27–30 (2000). doi:10.1093/nar/28.1.27.

[Kanehisa2019a] M. Kanehisa, "Toward understanding the origin and evolution of cellular organisms", *Protein Science* 28, 1947–1951 (2019). doi:10.1002/pro.3715.

[Kanehisa2021a] M. Kanehisa, M. Furumichi, Y. Sato, M. Ishiguro-Watanabe, and M. Tanabe, "KEGG: integrating viruses and cellular organisms", *Nucleic Acids Research* 49, gkaa970 (2020). doi:10.1093/nar/gkaa970.

[LiJJ2026a] J.-J. Li, H.-J. Wang, C. Zhang, Y. Li, J. Jiao, L.-L. Wang, C. Wu, D.-C. Zhong, Z.-Y. Wu, Z.-Y. Yu, and T.-B. Lu, "A high-flux membrane electrode assembly for $CO_2$ electroreduction to 4.5 M formate with over 8,000 h stability", *Nature Catalysis* 9, 492–501 (2026). doi:10.1038/s41929-026-01524-9.

[Liu2016a] C. Liu, B.C. Colón, M. Ziesack, P.A. Silver, and D.G. Nocera, "Water splitting biosynthetic system with $CO_2$ reduction efficiencies exceeding photosynthesis", *Science* 352, 1210–1213 (2016). doi:10.1126/science.aaf5039.

[Liu2024a] F. Liu, K. Miyatake, M. Tanabe, A.M.A. Mahmoud, V. Yadav, L. Guo, C.Y. Wong, F. Xian, T. Iwataki, M. Uchida, and K. Kakinuma, "High-Performance Anion Exchange Membrane Water Electrolyzers Enabled by Highly Gas Permeable and Dimensionally Stable Anion Exchange Ionomers", *Advanced Science* 11, e2402969 (2024). doi:10.1002/advs.202402969.

[Marecos2022b] S. Marecos, R. Brigham, A. Dressel, L. Gaul, L. Li, K. Satish, I. Tjokorda, J. Zheng, A.M. Schmitz, and B. Barstow, "Practical and thermodynamic constraints on electromicrobially accelerated $CO_2$ mineralization", *iScience* 25, 104769 (2022). doi:10.1016/j.isci.2022.104769.

[Park2021a] Y.S. Park, J. Jeong, Y. Noh, M.J. Jang, J. Lee, K.H. Lee, D.C. Lim, M.H. Seo, W.B. Kim, J. Yang, and S.M. Choi, "Commercial anion exchange membrane water electrolyzer stack through non-precious metal electrocatalysts", *Applied Catalysis B: Environmental* 292 (2021). doi:10.1016/j.apcatb.2021.120170.

[Ramdin2019a] M. Ramdin, A.R.T. Morrison, M. de Groen, R. van Haperen, R. de Kler, L.J.P. van den Broeke, J.P.M. Trusler, W. de Jong, and T.J.H. Vlugt, "High Pressure Electrochemical Reduction of $CO_2$ to Formic Acid/Formate: A Comparison between Bipolar Membranes and Cation Exchange Membranes", *Industrial & Engineering Chemistry Research* 58, 1834–1847 (2019). doi:10.1021/acs.iecr.8b04944.

[Riegraf2025a] M. Riegraf, M. Riedel, S.H. Jensen, S. Santhanam, S.A. Ansar, and M. Heddrich, "Solid Oxide Electrolysis Cells: Bridging Materials Development and Process System Engineering for Gigawatt-Scale Applications", *arXiv* (2025). doi:10.48550/arXiv.2512.16488.

[Salimijazi2020b] F. Salimijazi, J. Kim, A.M. Schmitz, R. Grenville, A. Bocarsly, and B. Barstow, "Constraints on the Efficiency of Engineered Electromicrobial Production", *Joule* 4, 2101–2130 (2020). doi:10.1016/j.joule.2020.08.010.

[Sheppard2023b] T.J. Sheppard, D.A. Specht, and B. Barstow, "Upper limit efficiency estimates for electromicrobial production of drop-in jet fuels", *Bioelectrochemistry* 154, 108506 (2023). doi:10.1016/j.bioelechem.2023.108506.

[Sheppard2024a] T.J. Sheppard, D.A. Specht, and B. Barstow, "Efficiency estimates for electromicrobial production of branched-chain hydrocarbons", *iScience* 27 (2024). doi:10.1016/j.isci.2023.108773.

[Siegel2015a] J.B. Siegel, A.L. Smith, S. Poust, A.J. Wargacki, A. Bar-Even, C. Louw, B.W. Shen, C.B. Eiben, H.M. Tran, E. Noor, J.L. Gallaher, J. Bale, Y. Yoshikuni, M.H. Gelb, J.D. Keasling, B.L. Stoddard, M.E. Lidstrom, and D. Baker, "Computational protein design enables a novel one-carbon assimilation pathway", *Proceedings of the National Academy of Sciences* 112, 3704–3709 (2015). doi:10.1073/pnas.1500545112.

[Siemens2019a] "Electricity-based fuels as a link between the electricity and transport sectors", Siemens AG (2019).

[Smolinka2011a] T. Smolinka, M. Günther, and J. Garche, "Stand und Entwicklungspotenzial der Wasserelektrolyse zur Herstellung von Wasserstoff aus regenerativen Energien", Fraunhofer ISE (2011).

[Torella2015a] J.P. Torella, C.J. Gagliardi, J.S. Chen, D.K. Bediako, B. Colón, J.C. Way, P.A. Silver, and D.G. Nocera, "Efficient solar-to-fuels production from a hybrid microbial-water-splitting catalyst system", *Proceedings of the National Academy of Sciences* 112, 2337–2342 (2015). doi:10.1073/pnas.1503606112.

[White2014a] J.L. White, J.T. Herb, J.J. Kaczur, P.W. Majsztrik, and A.B. Bocarsly, “Photons to formate: Efficient electrochemical solar energy conversion via reduction of carbon dioxide”, *Journal of CO2 Utilization* 7, 1–5 (2014). doi:10.1016/j.jcou.2014.05.002.

[Wise2022a] L. Wise, S. Marecos, K. Randolph, M. Hassan, E. Nshimyumukiza, J. Strouse, F. Salimijazi, and B. Barstow, “Thermodynamic Constraints on Electromicrobial Protein Production”, *Frontiers in Bioengineering and Biotechnology* 10 (2022). doi:10.3389/fbioe.2022.820384.

[Yang2017a] H. Yang, J.J. Kaczur, S.D. Sajjad, and R.I. Masel, “Electrochemical conversion of $CO_2$ to formic acid utilizing Sustainion™ membranes”, *Journal of CO2 Utilization* 20, 208–217 (2017). doi:10.1016/j.jcou.2017.04.011.

[YangH2020a] H. Yang, J.J. Kaczur, S.D. Sajjad, and R.I. Masel, “Performance and long-term stability of $CO_2$ conversion to formic acid using a three-compartment electrolyzer design”, *Journal of CO2 Utilization* 42 (2020). doi:10.1016/j.jcou.2020.101349.

[Ye2025a] N. Ye, K. Wang, Y. Tan, Z. Qian, H. Guo, C. Shang, Z. Lin, Q. Huang, Y. Liu, L. Li, Y. Gu, Y. Han, C. Zhou, M. Luo, and S. Guo, “Industrial-level $CO_2$ to formate conversion on Turing-structured electrocatalysts”, *Nature Synthesis* 4, 799–807 (2025). doi:10.1038/s44160-025-00769-9.

[Yu2024a] X. Yu, Y. Xu, L. Li, M. Zhang, W. Qin, F. Che, and M. Zhong, “Coverage enhancement accelerates acidic $CO_2$ electrolysis at ampere-level current with high energy and carbon efficiencies”, *Nature Communications* 15, 1711 (2024). doi:10.1038/s41467-024-45988-4.

[Yuan2022a] Y. Yuan, Q. Wang, Y. Qiao, X. Chen, Z. Yang, W. Lai, T. Chen, G. Zhang, H. Duan, M. Liu, and H. Huang, “*In Situ* Structural Reconstruction to Generate the Active Sites for $CO_2$ Electroreduction on Bismuth Ultrathin Nanosheets”, *Advanced Energy Materials* 12 (2022). doi:10.1002/aenm.202200970.

[Zeng2010a] K. Zeng and D. Zhang, “Recent progress in alkaline water electrolysis for hydrogen production and applications”, *Progress in Energy and Combustion Science* 36, 307–326 (2010).

[ZhangH2025a] H. Zhang, Z. Bo, M. Wang, W. Lin, Y. Liu, N. Liu, X. Sang, B. Yang, Z. Li, L. Lei, L. Dai, and Y. Hou, “Copper-stabilized bismuth subcarbonate electrocatalysts for durable large-scale formate production at kilowatt power”, *Nature Communications* 17, 577 (2025). doi:10.1038/s41467-025-67274-7.

[Zheng2025a] Y. Zheng, W. Ma, A. Serban, A. Allushi, and X. Hu, “Anion Exchange Membrane Water Electrolysis at 10 A $cm^{-2}$ Over 800 Hours”, *Angewandte Chemie International Edition* 64, e202413698 (2025). doi:10.1002/anie.202413698.
